# Core material and surface chemistry of Layer-by-Layer (LbL) nanoparticles independently direct uptake, transport, and trafficking in preclinical blood-brain barrier (BBB) models

**DOI:** 10.1101/2022.10.31.514595

**Authors:** Nicholas G. Lamson, Andrew J. Pickering, Jeffrey Wyckoff, Priya Ganesh, Joelle P. Straehla, Paula T. Hammond

**Affiliations:** Koch Institute for Integrative Cancer Research, Massachusetts Institute of Technology, Cambridge, MA 02142, USA; Department of Chemical Engineering, Massachusetts Institute of Technology, Cambridge, MA, 02142, USA; Department of Materials Science and Engineering, Massachusetts Institute of Technology, Cambridge, MA, 02142, USA; Department of Pediatric Oncology, Dana-Farber Cancer Institute, Boston, MA 02115, USA; Division of Pediatric Hematology/Oncology, Boston Children’s Hospital, Boston, MA 02115, USA; Broad Institute of MIT and Harvard, Cambridge, MA 02142, USA; Institute for Soldier Nanotechnologies, Massachusetts Institute of Technology, Cambridge, MA 02139, USA

**Keywords:** Blood-brain barrier, drug delivery, Layer-by-Layer, nanoparticles, intracellular trafficking, intravital imaging

## Abstract

Development of new treatments for neurological disorders, especially brain tumors and neurodegenerative diseases, is hampered by poor accumulation of new therapeutic candidates in the brain. Drug carrying nanoparticles are a promising strategy to deliver therapeutics, but there is a major need to understand interactions between nanomaterials and the cells of the blood-brain barrier (BBB), and to what degree these interactions can be predicted by preclinical models. Here, we use a library of eighteen layer-by-layer electrostatically assembled nanoparticles (LbL-NPs) to independently assess the impact of nanoparticle core stiffness and surface chemistry on *in vitro* uptake and transport in three common assays, as well as intracellular trafficking in hCMEC/D3 endothelial cells. We demonstrate that nanoparticle core stiffness impacts the magnitude of material transported, while surface chemistry influences how the nanoparticles are trafficked within the cell. Finally, we demonstrate that these factors similarly dictate *in vivo* BBB transport using intravital imaging through cranial windows in mice, and we discover that a hyaluronic acid surface chemistry provides an unpredicted boost to transport. Taken together, these findings highlight the importance of considering factors such as assay geometry, nanomaterial labelling strategies, and fluid flow in designing preclinical assays to improve nanoparticle screening throughput for drug delivery to the brain.

## Introduction

Neurological disorders, including but not limited to brain tumors and neurological diseases, are the leading global cause of years of life lost, and the second leading cause of death^1^. However, development of new treatments for neurologic disorders has been challenging due to poor drug transport across the blood-brain barrier (BBB) – the specialized vascular lining of the central nervous system^2^. As a result, at least 95% of newly discovered candidate therapies are excluded from entering the brain^3^, and there is a critical need to develop drug carriers that can transport therapeutic cargoes across the BBB.

Capillaries are the smallest blood vessels in the brain and are lined with endothelial cells supported by pericytes and astrocytes (**Figure 1a**). The endothelial cells comprise the majority of the mass transport barrier, with tight junctions stitching the cells together to exclude passive and non-specific transport between the cells. Uptake of nutrients, therapeutics, and drug carriers must rely on cell-mediated, active uptake to cross the BBB^4^. For this reason, the early stages of brain therapeutic development rely on *in vitro* BBB models that properly recapitulate endothelial cell morphology and transport mechanisms to screen candidates for brain delivery, necessitating standardized and high-throughput *in vitro* models to facilitate development of new delivery vehicles.

**Figure 1:**
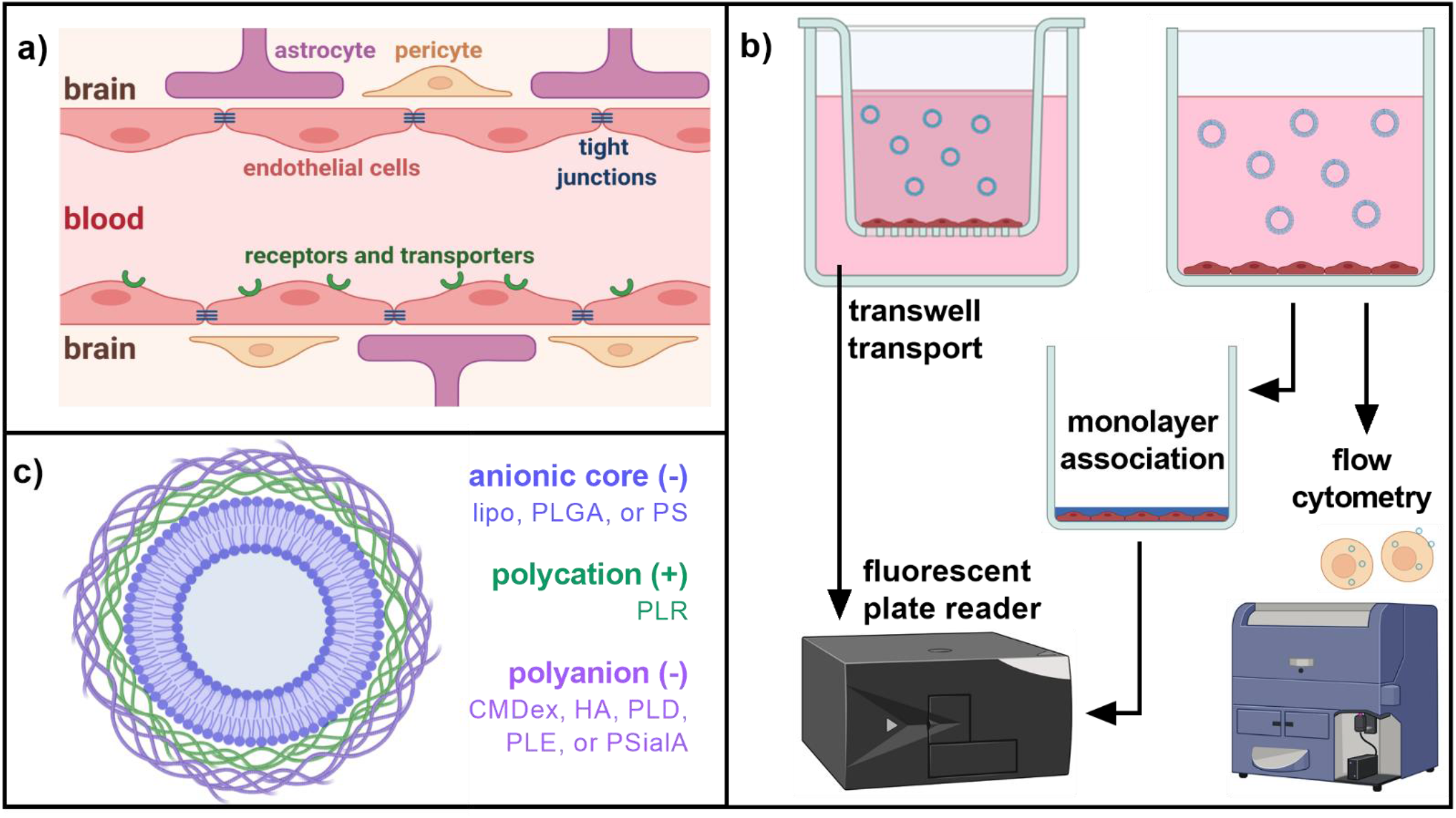
Probing nanomaterial uptake behavior in the blood brain barrier (BBB) using fluorescently labelled layer-by-layer nanoparticles (LbL-NPs). **(a)** The BBB derives its barrier properties primarily from endothelial cells, which are stitched together with tight junction proteins, but express receptors, transporters, and other machinery for cell-mediated uptake. **(b)** Using brain endothelial cell lines, three common metrics of nanomaterial uptake for *in vitro* BBB models are transwell transport, monolayer association, and flow cytometry using fluorescently tagged nanoparticles. The output for transwell transport is fluorescence of material that has passed through the cell layer at a given time, while monolayer association and flow cytometry respectively report fluorescence of cell populations or of individual cells. **(c)** LbL-NPs - used here to cross-compare the effects of nanoparticle stiffness and surface chemistry in BBB uptake models - begin with a fluorescently labelled and negatively charged core: anionic liposomes (lipo), acid-terminated poly (lactic-co-glycolic acid) (PLGA), or carboxylated polystyrene (PS). The cores are electrostatically complexed with a layer of polycation, in this case poly-L-arginine (PLR). Finally, a polyanion outer surface layer is added. The outer layers used here are carboxymethyldextran (CMDex), hyaluronic acid (HA), poly-L-aspartic acid (PLD), poly-L-glutamic acid (PLE), and polysialic acid (PSialA).

Within the field of *in vitro* BBB modelling, there are several available endothelial cell types, each of which present their own benefits and drawbacks. Primary brain endothelial cells, isolated from patient samples, are the most biologically sophisticated, but they lose their brain-specific endothelial phenotypes quickly in cell culture^5^. Endothelial cells derived from pluripotent stem cells are biologically diverse^6^, but these models’ recapitulation of the correct phenotype remains controversial^7^. Finally, immortalized cell lines are less genetically diverse, but reliably and stably express canonical BBB genes encoding junctional and transport proteins^8,9^. For this reason, we used the human-derived hCMEC/D3 immortalized microcapillary endothelial cell line. hCMEC/D3 has an endothelial phenotype, characteristic BBB protein expression^10–14^, and is known to recapitulate the endo-lysosomal and transcytosis systems of primary brain endothelial cells^15^.

Beyond type of endothelial cell used, there are also several model geometries – 2 or 3 dimensional, and with or without media flow – available. Each combination of these factors offers its own advantages for inter-lab consistency, ease of handling, and/or recapitulation of the physical environment of the brain capillaries^16^. For example, there are several 3D microfluidic models currently under development for screening of therapeutics or material libraries^17–19^. However, these remain difficult to standardize across labs and to scale for high-throughput studies. For these reasons, we have chosen to focus on validating more broadly accessible, 2D static models.

In addition to varying cell line and model geometry, and despite ample literature describing small molecule and biologic transport in BBB models^20,21^, there has been limited study of and little standardization in methodology for assessing BBB transport of nanomaterials^17,22^. Three commonly used assays include nanoparticle (NP) association via flow cytometry^23,24^, monolayer association of NPs^22,25^, and transport of NPs across transwell-grown cell monolayers^23,25^ (**Figure 1b**). Within the transwell model, there is not consensus for how transwell pore size or monolayer development time affect transport properties and gene expression^12,14,25^. By investigating these, we aim to better inform future, high-throughput materials screens to speed development of new strategies for drug delivery to the brain.

Here, we begin by probing relationships between hCMEC/D3 model growth parameters, barrier properties, and gene expression. Next, we employ a curated library of layer-by-layer assembled nanoparticles (LbL-NPs) to examine how these BBB models extend to nanomaterial uptake and transport behavior at the BBB. Our LbL-NP library combines three negatively charged NP cores – liposomes, poly(lactic-co-glycolic acid) (PLGA), and polystyrene (PS) – alone or with one of five polyanion outer layers – carboxymethyldextran (CMDex), hyaluronic acid (HA), poly-L-aspartic acid (PLD), poly-L-glutamic acid (PLE), or polysialic acid (PSialA) – to yield 18 formulations with defined core stiffness and surface chemistries (**Figure 1c**). Screening these NPs using the three previously described *in vitro* assays allows us to investigate how particle characteristics act as controlling factors for BBB uptake and transport, as well as how surface chemistry in particular dictates NP trafficking within hCMEC/D3 cells. Finally, to match the standard preclinical pipeline for therapeutic development, we use a select set of LbL-NPs to compare our *in vitro* BBB models to *in vivo* BBB transport in mice. We apply intravital multiphoton imaging through cranial windows to visualize NP concentrations in blood vessels and brain parenchyma, enabling us to calculate the NP permeability across the BBB^19,26^. By comparing these results to the *in vitro* assays, we conclude by highlighting promises and pitfalls of each method and how these can inform design of future studies for drug delivery to the brain.

## Results and Discussion

We began by characterizing the development of hCMEC/D3 endothelial cell monolayers on transwell membranes, to understand how choice of filter and duration of monolayer development affect the transport model. First, we seeded cells onto four different types of transwell filters – 0.4 μm pore polyester or polycarbonate, 1 μm pore polyester, or 3 μm pore polycarbonate – all coated with type 1 collagen to provide a suitable attachment and growing surface for the cells^10^. By trans-endothelial electrical resistance (TEER), a surrogate measure for barrier function of the monolayers^27^, cells on all four substrates formed a stable barrier within 48 hours (**Figure 2a**). This is consistent with other published studies^9^, as is a more gradual increase in barrier strength over at least four to seven days^11,14^. While all four filter types yielded similar TEER values, we noted that media leakage through the 3 μm pore polycarbonate during handling complicated experimental procedures. With that, and with concern that the smallest (0.4 μm) pores may provide non-negligible diffusive resistance to 100-200 nm nanomaterials, we chose to use the 1 μm pore polyester transwell filters for further investigation.

**Figure 2:**
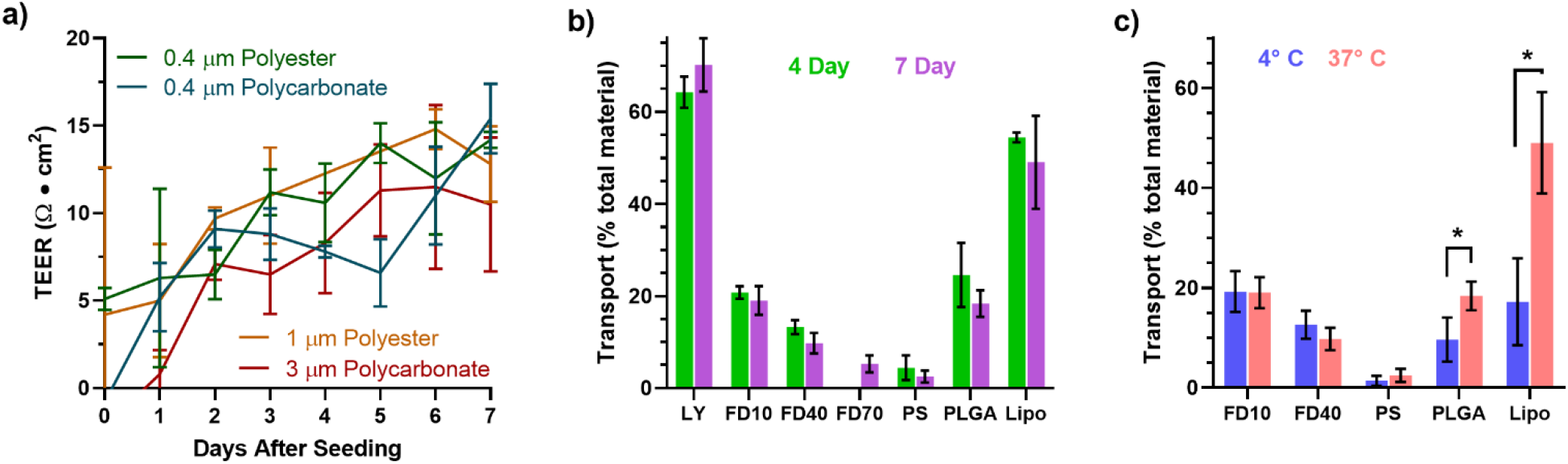
hCMEC/D3 cells grown on transwell inserts with 1 μm pores produce monolayers with proper barrier integrity to examine active transport of nanomaterials. (a) Across three pore sizes and two transwell membrane materials, transendothelial electrical resistance (TEER) of hCMEC/D3 monolayers initially climbed, then plateaued at three to five days post seeding. **(b)** On 1 μm pore polyester membranes, transport of paracellular permeability markers lucifer yellow (LY), 10 kDa FITC-dextran (FD10), 40 kDa FITC Dextran (FD40), and nanoparticle cores did not differ between monolayers grown for four or seven days. FD70 was examined on seven-day monolayers only. **(c)** In seven-day monolayers, performing transport studies at 4°C did not substantially diminish passive transport of diffusion markers FD10 and FD40. In contrast, PLGA nanoparticle and liposome transport slowed substantially, indicating that their uptake across monolayers is a predominantly active, transcellular process. Data display mean ± standard deviation for n = 3 transwell insert replicates (for TEER) or n = 3 biological plate replicates (for material transport). * p < 0.05 by Mann-Whitney U Test for non-parametric data.

We next investigated the barrier behavior of transwell monolayers at four or seven days past cell seeding, using passive diffusion markers to assess permeability of material through tight junctions between cells. As expected, smaller markers – Lucifer Yellow (LY) and 10 kDa FITC-Dextran (FD10) –are more permeable than larger substrates – 40 kDa FITC-Dextran (FD40) and 70 kDa FITC-Dextran (FD70) after a 24-hour incubation period (**Figure 2b**)^8^. There were no significant differences in any permeability between four-day and seven-day monolayers, indicating that model development time does not substantially affect these metrics. However, rapid passage of LY in particular corroborates previous reports that hCMEC/D3 monolayers do not recapitulate proper BBB exclusion of hydrophilic small molecules, but do provide a proper barrier to macromolecules and nanomaterials^28^. For anionic, 100 nm PS, PLGA, and liposomal NPs, there were also no significant transport differences between differing lengths of development. Notably, the substantially higher transport of PLGA NPs and liposomes than fluorescent dextrans – which would encounter less diffusive resistance based on their smaller sizes – indicates that these materials may cross the monolayers via an active process, rather than passively diffusing between the cells. To validate our hypothesis that PLGA and liposome transport is driven by cell-mediated, energy-dependent processes, we compared our standard seven-day monolayers to monolayers that were chilled to 4°C immediately after treatment addition (**Figure 2c**). While the passive diffusion markers did not demonstrate significant changes in permeability, substantially less of the PLGA NPs and liposomes crossed the cell monolayers at 4°C, confirming the transport to be active.

Having demonstrated similar barrier properties between transwell monolayers across culture conditions, we next investigated expression of endothelial genes and canonical BBB transporters. By preparation of complimentary DNA (cDNA) libraries and subsequent analysis via quantitative polymerase chain reaction (qPCR), we found no significant differences in seven genes – transferrin receptor (TFRC)^8^, low density lipoprotein related receptor 1 (LRP1)^12^, zonula occludens 1 (ZO-1 or TJP1)^5^, VE-cadherin (CDH5)^9^, CD34^29^, Von Willebrand Factor (VWF)^28^, and PECAM-1 (CD31)^28^ – across hCMEC/D3 monolayers developed for four, seven, or fourteen days (**Figure 3a**). Relative transcript levels matched closely with *in vitro* BBB studies across the literature^14,30^, and are tabulated in **Supplementary Table 1**. To extend these findings, we examined gene expression in cell monolayers grown on collagen-coated, non-porous plasticware for the same durations of time (**Figure 3b**). For the transferrin receptor, expression in transwell-grown and plasticware-grown cells matched well. For the remaining genes, however, there was substantially higher expression of the classical endothelial markers in transwell-grown cells on day four, though this abated by day seven. Therefore, to draw the most appropriate comparisons between the two assay formats, we decided to move forward with seven-day monolayers for the remainder of our experiments. By immunofluorescence confocal imaging, we confirmed that monolayers grown in both systems were structurally similar, with expected patterns of localization for ZO-1 (**Figures 3c,d**), Cadherin-5 (**Figures 3e,f**), and coordinated F-actin fibers across multiple cells (**Figure 3g,h**). The discontinuous nature of ZO-1 and cadherin 5 around each cell, in particular, is consistent with previous findings and supports reports that hCMEC/D3 monolayers are leaky to small molecules, but can provide sufficient transport barriers for macromolecules and nanomaterials^9,10,28,31^.

**Figure 3:**
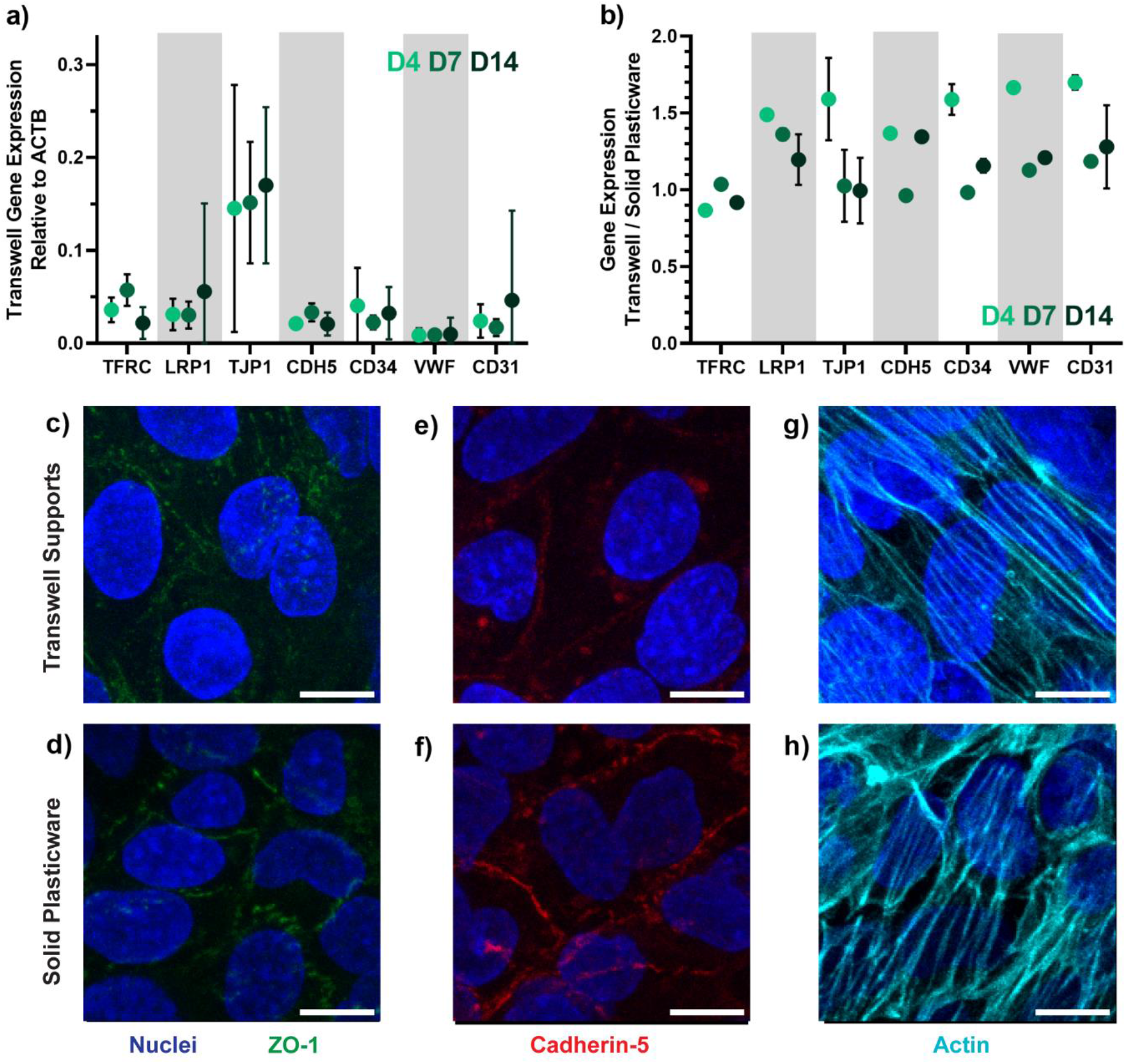
hCMEC/D3 cells demonstrate consistent gene expression and localization of key cellular proteins across experimental conditions. **(a)** Gene expression via by RT-qPCR is similar over time, comparing cells grown on transwell filters for 4, 7, or 14 days. For each gene, there was no significant difference between groups by Kruskal–Wallis *H* test, and error bars display standard deviation of three biological replicates at passage numbers 2, 6, and 10. **(b)** Likewise, the difference in gene expression did not vary substantially between cell monolayers grown on transwell filters and those grown on solid plasticware. **(c)** By confocal imaging, ZO-1 in both transwell-grown cells and **(d)** plasticware-grown cells show discontinues rings of tight junctions around each cell. **(d)** The same pattern is seen in cadherin-5 in both transwell-grown and **(f)** plasticware-grown cells. **(g)** Actin staining in transwell-grown and **(h)** plasticware-grown cells shows cooperative filament organization across several cells. Scale bars display 10 μm.

Knowing that transwell-grown and plasticware-grown cells did not differ in key gene expression, we next created a library of LbL-NPs to use in cell assays. The modularity of LbL-NPs allows for combinatorial screening of NP core materials and surface chemistries, to assess the independent effects of changing each factor on their biological activity^32^. For our library, anionic (phospholipid) liposomes, acid-terminated PLGA NPs, or carboxylated PS NPs with diameters of 80-100 nm and covalently bound fluorophores were first layered with a coating of poly-L-arginine (PLR) to achieve zeta potential charge conversion to > 40 mV. The PLR-coated NPs were then layered with one of five polyanions – carboxymethyldextran (CMDex), hyaluronic acid (HA), poly-L-aspartic acid (PLD), poly-L-glutamic acid (PLE), or polysialic acid (PSialA) – chosen to give anionic outer surfaces with zeta potential < −30 mV, as well as cover a range of synthetic polypeptides, synthetic carbohydrates, and naturally occurring carbohydrates. Characterization by dynamic light scattering (DLS) is displayed in **Table 1**.

**Table 1:** NP characteristics measured by dynamic light scattering. All values are displayed as mean ± SD of three runs. PDI: polydispersity index.

| **Table 1:** NP characteristics measured by dynamic light scattering. All values are displayed as mean ± SD of three runs. PDI: polydispersity index. | | | | |
| --- | --- | --- | --- | --- |
| NP Name | Number Average Size (nm) | Z-Average Size (nm) | PDI | Zeta Potential (mV) |
| Bare Liposome | 82.9 ± 5.9 | 105.2 ± 1.5 | 0.11 ± 0.03 | -52.2 ± 0.9 |
| Lipo-PLR-CMDex | 102.1 ± 13.8 | 167.8 ± 4.4 | 0.23 ± 0.09 | -34.8 ± 1.0 |
| Lipo-PLR-HA | 88.2 ± 7.5 | 124.9 ± 2.1 | 0.20 ± 0.01 | -33.0 ± 1.1 |
| Lipo-PLR-PLD | 81.6 ± 5.6 | 131.9 ± 4.7 | 0.22 ± 0.02 | -42.9 ± 7.2 |
| Lipo-PLR-PLE | 82.6 ± 8.5 | 130.2 ± 2.6 | 0.21 ± 0.01 | -49.0 ± 3.8 |
| Lipo-PLR-PSialA | 104.2 ± 3.1 | 144.6 ± 3.4 | 0.11 ± 0.03 | -50.0 ± 2.7 |
| Bare PLGA | 85.0 ± 4.9 | 136.7 ± 3.1 | 0.17 ± 0.02 | -51.0 ± 2.3 |
| PLGA-PLR-CMDex | 85.0 ± 4.9 | 167.8 ± 4.4 | 0.17 ± 0.02 | -51.0 ± 2.3 |
| PLGA-PLR-HA | 88.5 ± 3.8 | 144.3 ± 3.9 | 0.25 ± 0.05 | -30.5 ± 0.3 |
| PLGA-PLR-PLD | 75.8 ± 8.7 | 148.9 ± 5.5 | 0.30 ± 0.05 | -41.8 ± 1.3 |
| PLGA-PLR-PLE | 72.7 ± 10.8 | 215.9 ± 9.0 | 0.17 ± 0.04 | -42.6 ± 0.5 |
| PLGA-PLR-PSialA | 78.7 ± 10.2 | 152.2 ± 1.9 | 0.36 ± 0.04 | -43.5 ± 3.7 |
| Bare PS | 107.7 ± 7.0 | 131.4 ± 2.3 | 0.07 ± 0.05 | -59.0 ± 0.6 |
| PS-PLR-CMDex | 148.5 ± 11.1 | 143.2 ± 1.1 | 0.11 ± 0.08 | -52.5 ± 1.8 |
| PS-PLR-HA | 98.7 ± 6.7 | 120.8 ± 24.5 | 0.18 ± 0.05 | -31.9 ± 1.0 |
| PS-PLR-PLD | 108.6 ± 7.4 | 143.1 ± 4.5 | 0.08 ± 0.05 | -43.7 ± 0.6 |
| PS-PLR-PLE | 99.4 ± 10.2 | 166.0 ± 3.7 | 0.28 ± 0.04 | -46.7 ± 2.3 |
| PS-PLR-PSialA | 109.2 ± 2.4 | 133.2 ± 1.1 | 0.01 ± 0.00 | -54.0 ± 3.3 |

We applied our LbL-NP library to the three orthogonal uptake or transport assays using the hCMEC/D3 endothelial cells: flow cytometry and monolayer association using cells grown in standard plasticware, or transport across a transwell monolayer. In flow cytometry, differing fluorophores and/or brightness between the NP cores precludes direct comparison across the groups. Within their formulation groups, polystyrene core LbL-NPs (**Figure 4a**), PLGA-core LbL-NPs (**Figure 4b**), and liposome core LbL-NPs (**Figure 4c**) all displayed improvements in median fluorescence intensity (MFI) over the bare NP cores, indicating that LbL functionalization generally improved interactions with the hCMEC/D3 cells. For PS or PLGA core LbL-NPs, the polypeptide outer layers PLD and PLE conferred an uptake advantage, while liposome core LbL-NPs with polysaccharide outer layers CMDex and HA showed stronger uptake than other formulations. However, most of the groups showed only up to 2- to 3-fold difference between LbL-NP formulations, which is consistent with other flow cytometry studies of NP surface chemistry impacts on uptake in hCMEC/D3^23^, as well as in other cell and tissue types^33,34^.

**Figure 4:**
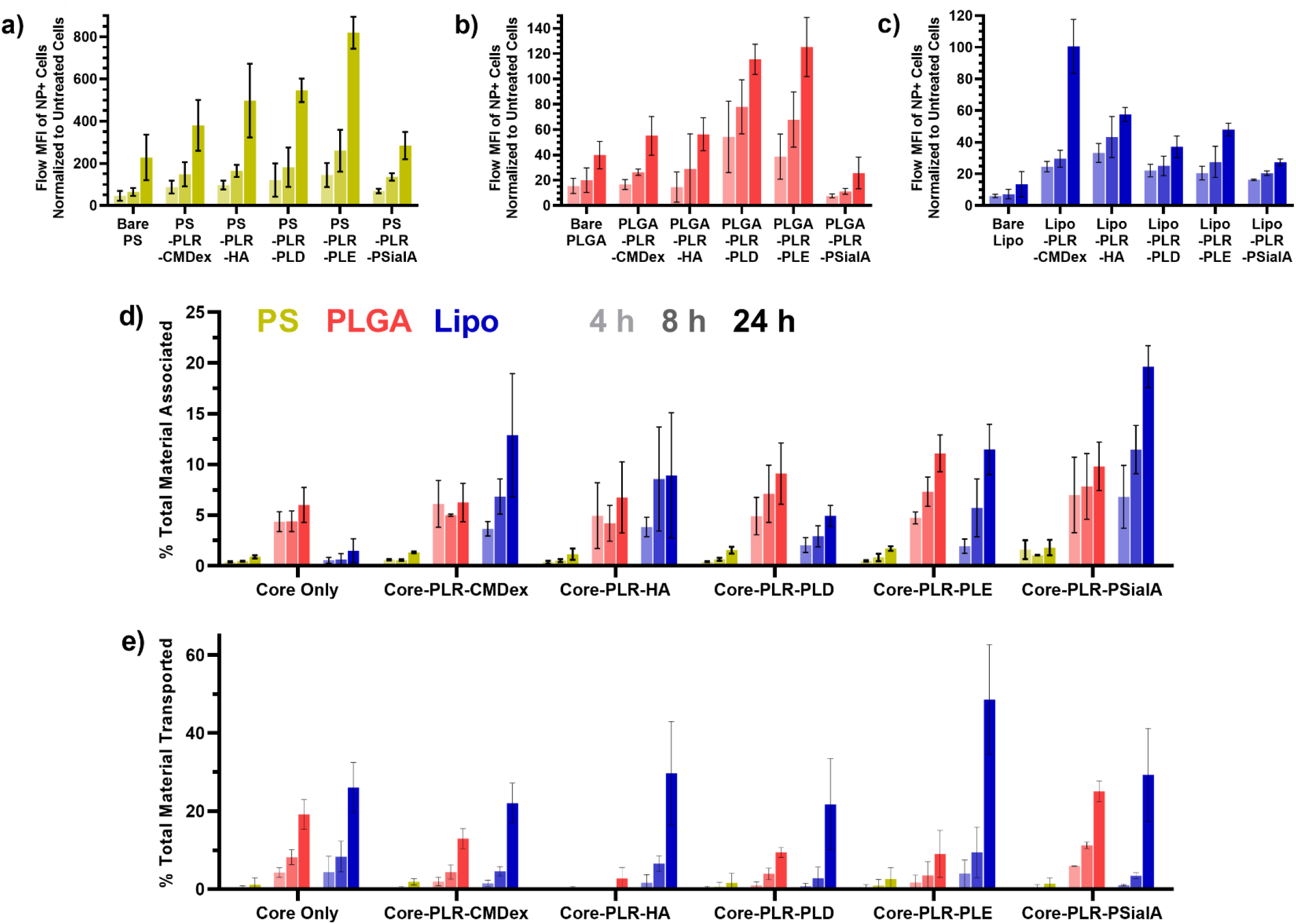
hCMEC/D3 based models of the BBB differ in their uptake of a library of layer-by-layer assembled nanoparticles. **(a)** In cells grown on solid plasticware and treated with PS core nanoparticles, flow cytometry mean fluorescence intensity for nanoparticle positive cells differed slightly by outer layer chemistry, with PLE outperforming the others. This trend was mirrored closely in **(b)** cells treated with PLGA core nanoparticles, while **(c)** cells treated with liposome-core nanoparticles showed greatest uptake with CMDex capped particles instead. **(d)** Monolayer association in cells grown on solid plasticware indicated a preference for soft nanoparticle uptake, with the exception of bare or PLD-coated liposomes. **(e)** Likewise, in monolayers grown on transwells, soft liposome-core nanoparticles transported to a greater degree than the stiffer polymer nanoparticles, regardless of surface chemistry. Data is reported as arithmetic mean ± standard deviation of three plate replicates, with three technical replicate wells per treatment per plate.

We next examined the LbL-NP library for uptake in monolayer association (**Figure 4d**) and for transport through monolayers grown on transwell supports (**Figure 4e**), both of which offer the advantage of direct cross-comparison between LbL-NP formulations with different NP cores. In both cases, uptake or transport of the liposome based LbL-NPs was greater than or similar to the stiffer PLGA NPs with matched outer surface layers, which substantially outperformed the PS core LbL-NPs with the same surface chemistries. As LbL films cover the surface of each core NP, we hypothesize that differences can be primarily attributed to the stiffness and deformability of the core material. This improved performance of softer nanomaterial transport is consistent with studies of nanogels in static hCMEC/D3 systems^35^, while limited studies in systems with fluid flow have not yet reached consensus on the relationship between particle stiffness and BBB transport^17,19,36^. By comparing surface chemistries, we once again saw that LbL functionalization improved monolayer association (**Figure 4d**) for most formulations, though the differences were less stark than those observed by flow cytometry. We did not observe this relationship for transwell transport (**Figure 4e**), in which bare PS and PLGA particles transported as well as, or better than, several of the polymer functionalized formulations.

While measuring NP fluorescence in monolayer association and transwell transport assays, we found that it was necessary to fully homogenize and solubilize our samples to obtain quantitative data. In the transwell assay, reading fluorescence directly from lower chamber implied 160-180 % of theoretically maximum NP transport for PLE coated liposomes (**Supplementary Figure 1a**). By homogenizing the samples – including calibration curves – with dimethyl sulfoxide (DMSO) to break up the liposomes and heparin sulfate to sequester PLR away from the sulfo-Cy5 fluorophore, we obtained the data displayed in **Figures 4d** and **4e**. Because homogenizing the particles led to a brighter but more consistent fluorescence (**Supplementary Figure 1b**), we concluded that the Cy5 fluorophore attached to the liposome core is partially self-quenching, and liposome breakup allows for higher activity of solubilized lipid-fluorophore conjugates. By contrast, PLGA core LbL-NPs, which do not self-quench, do not demonstrate data discrepancies (**Supplementary Figure 1c**). The inflated transport values of some liposomal formulations thus implied that they were degrading as they transported across the cell monolayers. This led us to hypothesize that while core identity has the largest impact on the total amount of NP transported, outer surface chemistry dictates the intracellular mechanisms by which particles are transported and processed.

To test this hypothesis, we selected three LbL-NP formulations (**Supplementary Table 2**) with liposome cores to examine their intracellular trafficking by confocal microscopy in transwell-grown monolayers: bare liposomes as a formulation undergoing no apparent degradation during transport, HA coated NPs as a formulation undergoing minimal degradation, and PLE coated NPs as a formulation undergoing massive degradation. After eight hours of NP treatment, we imaged z-stacks of the NP-treated monolayers to construct orthogonal views, and we observed that Cy5 signal for bare liposomes and HA-coated LbL-NPs presented as small spots scattered throughout the cell bodies (**Figure 5**). By contrast, the PLE-coated LbL-NPs demonstrated heavy localization of large, bright areas of NP signal to the apical surfaces of the cells. We confirmed that PLE NPs remain colloidally stable in cell culture media over this time period as measured by DLS (**Supplementary Figure 2**), and therefore this morphology is not likely related to be sedimenting at the cell surface, but rather indicates that PLE NPs traffic through different intracellular mechanisms than the bare liposomes or HA-coated LbL-NPs.

**Figure 5:**
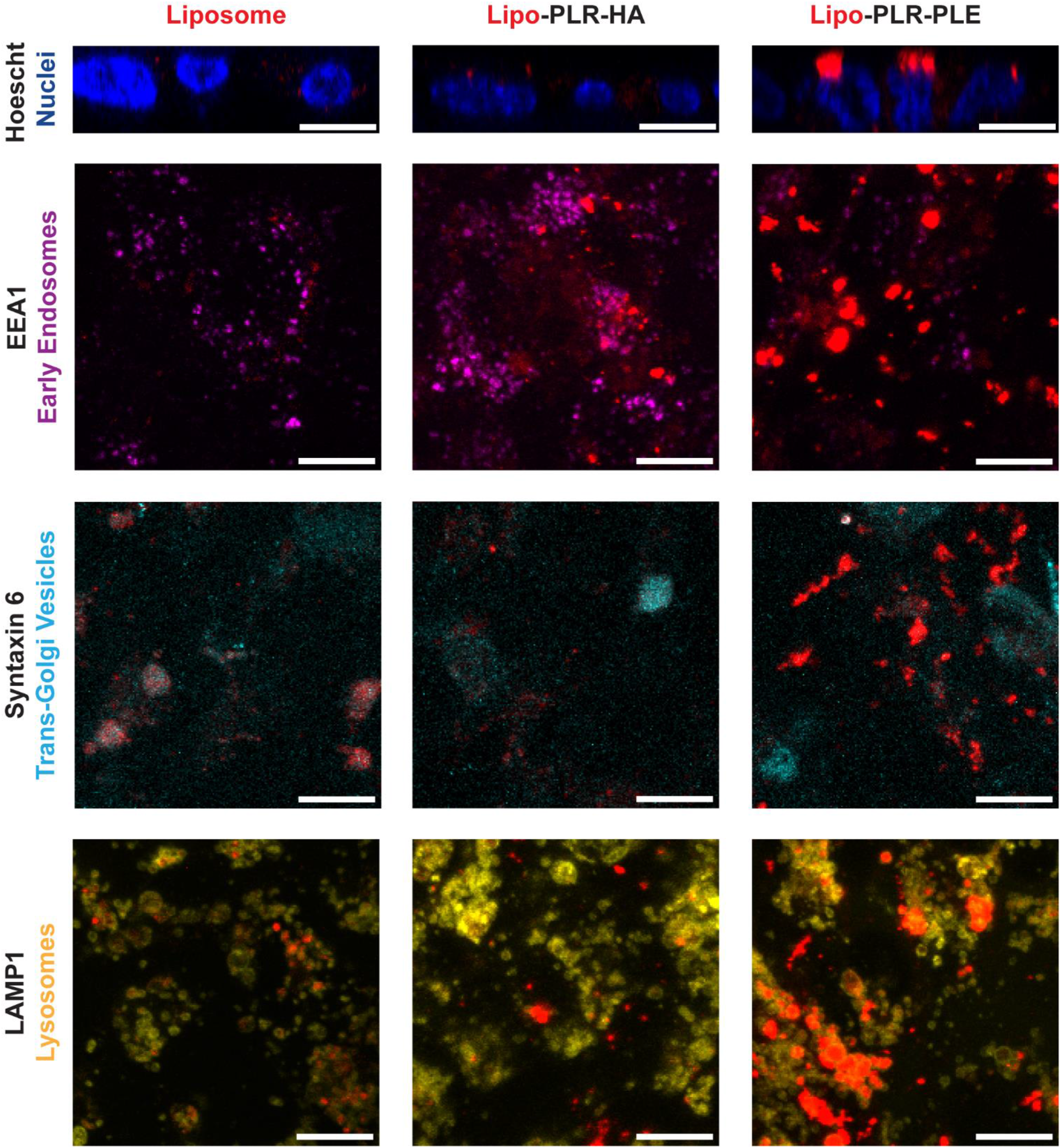
Outer layer chemistries on liposome-core LbL-NPs influence cellular uptake and trafficking in hCMEC/D3 monolayers grown on transwell supports. In orthogonal projections, bare liposomes and HA-coated LbL-NPs (Lipo-PLR-HA) can be seen distributing throughout the thickness of the monolayers, while PLE-coated NPs (Lipo-PLR-PLE) are concentrated in large aggregates near the apical surface. None of the nanoparticles demonstrated strong association with the EEA1 marker of early endosomes, suggesting that processing through this stage occurs quickly. Bare liposomes colocalized modestly with the Syntaxin 6, HA-NPs showed partial association, and PLE-NPs did not. LAMP1 staining was largely independent of NP signal for liposomes and HA-NPs, but aggregated PLE-NP signals appear to be enclosed in lysosomal vesicles. Scale bars display 10 μm. Bare liposomes required higher laser power for proper visualization, so quantity of nanoparticle signal should not be directly compared between particles.

To further examine intracellular trafficking, we co-stained identically treated cells with markers for several uptake and transport related proteins. Neither clathrin nor caveolin-1 showed any substantial overlap in signal with any of the NP particle formulations (**Supplementary Figure 3**), suggesting that all three are taken up into cells by the same macropinocytosis pathway. Similarly, none of the formulations visibly colocalized with the early endosome marker EEA1 (**Figure 5**), indicating that all of the NPs are processed quickly through early endosomes and into other intracellular compartments. We then investigated two major downstream pathways, and determined that bare liposomes and HA-coated LbL-NPs show strongest localization with Syntaxin 6, a marker for Golgi-associated vesicles that drive transcytosis^37^. By contrast, the PLE-coated LbL-NPs accumulate predominantly in vesicles with high expression of LAMP1, a lysosomal marker that denotes vesicles with high degradation capacity^38^. This difference in colocalization supports our hypothesis that the NP surface chemistry dictates intracellular trafficking, explains the inflated fluorescence data (Supplementary Figure 1a), and indicates that liposomes and HA-coated LbL-NPs transport through endothelial cells intact, while PLE-coated NPs are degraded during transport.

Having investigated the role of core stiffness and surface chemistry on uptake and transport of NPs by BBB endothelial cells *in vitro*, we finally sought to understand the extent to which these controlling factors are recapitulated *in vivo*. To do so, we used a select set of LbL-NPs (**Supplementary Table 2**) to examine for BBB permeability via intravital imaging through a cranial window in mice. This method has been used in previous studies to construct static images of NP accumulation in brain^39^, and has recently been adapted to capture a time series of 3-dimensional images for quantitative permeability measurements between the brain capillaries and parenchyma^19^. Briefly, fluorescent dextran and fluorescent NPs are administered systemically, then a cranial window is generated and two-photon microscopy is used to generate a time series of multiphoton confocal images. The dextran signal from the first imaging time point is converted to a 3-dimensional mask to differentiate blood vessels from brain parenchyma, and the subsequent images are compared to this mask to determine permeability of the NPs across the BBB (**Figure 6a)**. As a passive diffusion marker, the consistent and slow leakage of dextran out of the blood vessels was also computed to use as a normalization factor, eliminating effects of any slight Z-shifts over time that could not be accounted for by differential slice selection.

**Figure 6:**
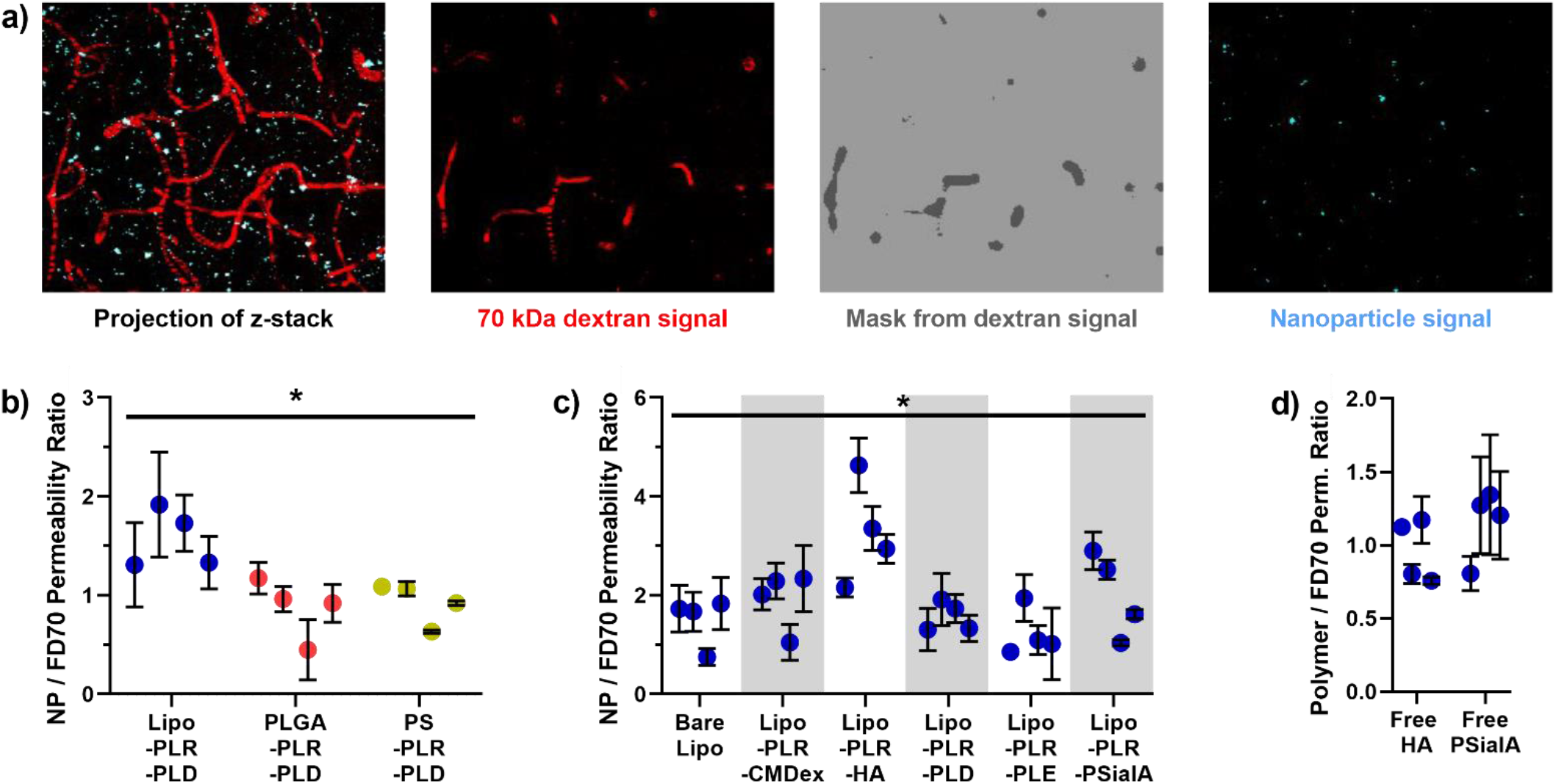
Intravital imaging enables the calculation of nanoparticle permeability across the blood brain barrier in mice. **(a)** From sequential z-stack images, the starting signal for a 70 kDa FITC-dextran (FD70) blood vessel marker is used to create a mask separating vessel pixels from brain parenchyma. This mask is applied to both dextran and nanoparticle signal for the first and all subsequent time points, allowing for calculations nanoparticle transport out of the vessels and into the brain. **(b)** For PLD outer layer nanoparticles with differing cores, liposome based materials displayed slightly higher BBB permeability than stiffer polymer particles. **(c)** For liposome core particles, an HA outer layer appeared to confer an advantage to crossing the blood brain barrier compared to other outer surface layers, while PSialA coated nanoparticles demonstrated high variability between mice. **(d)** However, size-matched HA and PSialA polymers did not cross the blood brain barrier faster than the biologically inert dextran. Data display mean ± standard error for individual animals, based on 3-15 permeability measurements, depending on length of experiment viability. ^*^ Treatment groups are statistically different (*p* < 0.005) by Kruskal-Wallis *H* test.

After validating that our method of retroorbital NP injection yielded comparable results to the previously published method of tail vein injections (**Figure S4**), we determined the BBB permeability of our selected LbL-NPs in mice, and all tabulated permeability values are available in **Supplementary Table 3**. For a constant PLD outer layer, softer liposome-based NPs had higher permeability than PLGA and PS core LbL-NPs (**Figure 6b**), although the difference was less stark than in the *in vitro* assays. Similarly, among liposome core LbL-NPs, varying surface chemistry generally had a modest impact on BBB permeability (**Figure 6c**), and LbL functionalization did not boost the transport of the NPs compared to the bare liposomes, consistent with observations from the transwell transport assay. Unexpectedly, we observed that HA-coated NPs significantly out-performed the other formulations, including the other carboxylated carbohydrates CMDex and PSialA. HA is a known binding partner for several receptors and proteins – especially CD44 – which are known to express in brain endothelial cells, including in the hCMEC/D3 cell line^30^; however, CD44 has not been implicated in any transcytotic or other uptake activity in endothelial cells^40^. We hypothesized that the particles were instead undergoing random binding events with endothelial cell surface proteins as they passed through the capillaries. This phenomenon would slow particles down in the blood flow environment and enhance their localization to the BBB surface, allowing for greater nonspecific uptake by macropinocytosis, as was implied from our *in vitro* microscopy analysis. To probe this hypothesis, we created Cy5-labelled HA and PSialA polymers, size matched to the FD70 diffusion marker, and examined their BBB permeability (**Figure 6d**). Because neither of these polymers transported faster than the passively diffusing dextran, we can conclude that the polymer interactions with cell surfaces do not, on their own, drive active uptake and transcytosis. While this supports our hypothesis that HA coatings improve NP permeability by changing their flow behavior at the BBB, future development of these NPs as promising drug delivery carriers will include further characterization in flow-based models to investigate interactions with specific binding partners on human and murine endothelial cells.

## Conclusions

Our data highlight major strengths and weaknesses for *in vitro* metrics of nanomaterial uptake and transport at the BBB. hCMEC/D3, the most commonly applied immortalized cell line for BBB modeling, expresses tight junctions that do not recapitulate BBB exclusion of small molecules, but do provide an appropriate barrier for studying interactions with nano-scale materials. We have demonstrated that cell assay behavior and gene expression do not change across a range of commonly used development times and cell culture substrates, allowing for cross-comparison across assays in the literature. For the three most common *in vitro* metrics of nanomaterial uptake and transport – flow cytometry, monolayer association, and transwell transport – flow cytometry provides the most comparable data to most NP uptake studies in other cell and tissue types. However, monolayer association and transwell transport allow for better cross-comparison across different NP cores in LbL-NP formulations. Specifically, the NP core consistently has the greatest effect on total uptake, with softer liposome cores enjoying the greatest magnitude of NP uptake or transport. In contrast, surface chemistry is a stronger determinant of intracellular trafficking patterns, and whether a NP is transported across the BBB intact, though transwell transport was the only assay that fully captured these differences. The relative importance of LbL-NP core stiffness and surface chemistries were conserved for permeability across the mouse BBB as measured via intravital imaging, though none of the *in vitro* models predicted HA functionalization to be the standout formulation *in vivo*. Taken together, these data underscore that, while transwell transport gives the most complete picture of NP behavior in BBB endothelial cells, *in vitro* uptake assays alone are not sufficient to identify top candidates for NP drug delivery carriers to the brain; instead, these metrics must be combined with careful mechanistic understanding of how the materials interact with cells, as well as judicious use of preclinical *in vivo* models, to provide comprehensive evidence for clinical translational potential.

## Acknowledgements

The authors would like to thank the Koch Institute’s Robert A. Swanson (1969) Biotechnology Center for technical support, specifically the Flow Cytometry Core, High Throughput Sciences Core, and Microscopy Core, as well as the MIT BioMicro Center. This work was supported in part by the Koch Institute Support (core) Grant P30-CA14051 from the National Cancer Institute. We would also like to acknowledge funding from Cancer Research UK and the Brain Tumour Charity grant REF: C42454/A28596. N.G.L. was supported by a postdoctoral fellowship from the Ludwig Center at MIT’s Koch Institute for Integrative Cancer Research, as well as the Convergence Scholars Program at MIT. A.J.P. was supported by the Natural Sciences and Engineering Research Council of Canada (NSERC), [CGSD3 - 557538 - 2021]. J.P.S was supported by the Charles W. (1955) and Jennifer C. Johnson Cancer Research Fund, Cannonball Kids’ cancer Foundation, and the Rally Foundation for Childhood Cancer Research. Figure 1 was created with BioRender.com. Finally, we would like to thank Dr. Tamara G. Dacoba for her help editing this manuscript.

## Plain Language Summary

Nanoparticles are a promising strategy to get new therapies into the brain. Here, we show that softer nanoparticles deliver more efficiently, and that surface chemistry dictates whether the particles enter the brain space fully intact or broken down into smaller components.

## Materials and Methods

### 1. Materials

1,2-distearoyl-sn-glycero-3-phospho-(1’-rac-glycerol) sodium salt (DSPG), 1,2-distearoyl-sn-glycero-3-phosphoethanolamine (DSPE), 1,2-distearoyl-sn-glycero-3-phosphocholine (DSPC) and cholesterol were purchased from Avanti. Sulfo-cyanine dyes with NHS ester or amine handles were purchased from Lumiprobe. Chloroform was purchased from TCI. Methanol, Poly(D,L-lactide-glycolide) (PLGA Resomer RG502H, 7-17 kDa), Rhodamine B – PLGA (50:50 monomer ratio, 10-30 kDa), the hCMEC/D3 cell line, Accumax dissociation reagent, Type 1 rat tail collagen, ascorbic acid, β-mercaptoethanol, lucifer yellow, FITC-labelled dextrans, dimethyl sulfoxide (DMSO), heparin sulfate, bovine serum albumin (BSA), saponin, and sulfo-N-hydroxysuccinimide were purchased from Millipore Sigma. Cy5-functionalized PLGA (10-15 kDa) was purchased from PolySciTech. Whatman Nucleopore polycarbonate hydrophilic membranes (400, 200, 100 and 50 nm sizes) were purchased from GE. 50/15 mL Falcon tubes, DNA LoBind tubes, 10% neutral buffered formalin, Polystyrene semi-micro cuvettes, 0.22 μm polyethersulfone syringe filters, Spectrapor dialysis membranes, microscope slides, coverslips, and slide sealer nail polish were purchased from VWR. D02-E300-05-ND, 02-E100-05-N, and C02-E100-05-N tangential flow filtration filters were purchased from Repligen. Poly-L-arginine hydrochloride (38.5 kDa), poly-L-aspartic acid (14 kDa), and poly-L-glutamic acid (15 kDa) were purchased from Alamanda Polymers. Hyaluronic acid (40 kDa) was purchased from LifeCore Biomedical. Carboxymehtyldextran and polysialic acid were purchased from Carbosynth. DTS 1070 folded capillary zeta cells were purchased from Malvern. Tissue culture plasticware (T75, T25, clear and white 96 well plates), 24-well 1 μm pore transwell plates, individual transwell inserts, Penicillin/Streptomycin and fetal bovine serum (FBS) were purchased from Corning. EMB-2 cell culture media was purchased from Lonza. Phosphate buffered saline (PBS), chemically defined lipid concentrate, LabTek 8-chamber coverslips, Hoeschst 33342, fluorescently labelled wheat germ agglutinin, fluorescently labelled phalloidin, AlexaFluor 488-labelled anti-VE Cadherin (16B1), AlexaFluor 488-labelled anti-ZO-1 (ZO1-1A12), 5 M bioreagent grade NaCl solution, 1 M bioreagent-grade HEPES, chemically defined lipid concentrate, basic fibroblast growth factor, PCR tube strips with caps, Pierce endotoxin removal columns, and yellow-green or red fluorescent polystyrene microspheres (100 nm Fluospheres) were purchased from Thermo Fisher. RNeasy Plus Mini Kits for RNA extraction and QuantiTect Reverse Transcriptase Kits were purchased from Qiagen. Roche Light Cycler-DNA Master SYBR Green I mastermix and Corning Axygen 384-well PCR microplates were purchased through the MIT BioMicro Center / KI Genomics Core. IDTE buffer, nuclease free water, and PrimeTime PCR Primers were purchased from Integrated DNA Technologies (IDT). The Voltohmmeter and accompanying electrodes were purchased from World Precision Instruments. 1-Ethyl-3-[3-dimethylaminopropyl]carbodiimide hydrochloride (EDC) was purchased from Chem-Impex. Falcon cell strainer tubes were purchased from Fisher Scientific. Syntaxin 6 (C34B2) Rabbit mAb 2869, Caveolin-1 (D46G3) XP^®^ Rabbit mAb 3267, Clathrin Heavy Chain (D3C6) XP® Rabbit mAb 4796, EEA1 (C45B10) Rabbit mAb 3288, and Anti-rabbit IgG (H+L), F(ab’)2 Fragment (Alexa Fluor® 488 Conjugate) #4412 were purchased from Cell Signaling Technologies. 6 mm biopsy punched were purchased from McKesson. VECTASHIELD Antifade Mounting Medium (H-1000) was purchased from Vector Laboratories.

### 2. Nanoparticle Synthesis

#### 2.1. Liposome Synthesis

Cholesterol and lipid stocks were made in chloroform and methanol, then combined in round bottom flask at a mol ratio of 31 Chol : 31 DSPC : 7 DSPE : 31 DSPG. The lipids were dried into a thin film using a BUCHI RotoVap system under heat (55 °C, water bath) until completely dry (<30 mBar). A Branson sonicator bath was heated to 65°C, at which point the RBF with the lipid film was partially submerged in the bath and a volume of Milli-Q deionized water was added to re-suspend the lipid film to a 1 mg lipid/mL solution. The liposome solution was sonicated three times for [1 minute on, 1 minute off], then transferred to an Avestin LiposoFast LF-50 liposome extruder. The extruder was connected to a Cole-Parmer Polystat Heated Recirculator Bath to maintain a temperature of 65 °C. The liposomes were extruded through nucleopore membranes until a 50-100 nm liposome was obtained. Typically, this was achieved by passing through stacked 400 and 200 nm membranes, 100 nm, and 50 nm membranes. The liposomes were analyzed by dynamic light scattering (DLS) to verify sizes less than 100 nm and PDI values less than 0.2. Fluorescently labeled liposomes were prepared via NHS-coupling of Sulfo-cyanine dyes with NHS ester handles to DSPE head group amines; reactions in 15 mM sodium carbonate buffer (pH 9), stirred at room temperature overnight. Unconjugated dye was purified away from the labelled liposomes by tangential flow filtration (TFF) before characterization of the liposomes by DLS.

#### 2.2. PLGA Nanoparticle Synthesis

10 mg PLGA polymers was dissolved at a concentration of 5 mg/ml in acetone, with a mass ratio of 9:1 Resomer 502H to dyed (RhodB or Cy5) polymer. 12 ml of Milli-Q water was added to a scintillation vial and stirred gently on a stir plate while heating to 35°C. The PLGA solution was drawn up in a syringe with a 26-gauge needle then slowly added to the water under constant stirring. An additional 8 mL of 35°C deionized water was added, and the vial was left to stir, uncovered under ventilation, for at least two hours. Particles were characterized by DLS and concentrated to 1 mg/mL by TFF.

#### 2.3. Layer-by-Layer Polymer Functionalization

Nanoparticles were layered by adding nanoparticle solution (0.5 mg/mL when layering HA, 1 mg/mL otherwise) to an equal volume of polyelectrolyte solution under sonication. The mixture was sonicated for approximately three seconds then vortexed for approximately ten seconds. The weight equivalents (wt. eq.) of polyelectrolyte used with respect to liposome core were 0.4 for poly-L-arginine (PLE), 1.6 for carboxymethyldextran (CMDex), 1.2 for hyaluronic acid (HA), 0.8 for poly-L-aspartic acid (PLD), 3 for poly-L-glutamic acid (PLE), and 1.4 for polysialic acid (PSialA). Polyelectrolyte solutions were prepared in 50 mM HEPES (pH 7.4) and 40 mM NaCl, with the exception of HA, which was prepared in 2 mM HEPES. The freshly layered particles were allowed to incubate at room temperature for 5 to 30 minutes, then purified using the tangential flow filtration.

#### 2.4. Tangential Flow Filtration

Nanoparticle samples to be purified were connected to a Spectrum Labs KrosFlo II filtration system using Masterflex, Teflon coated tubing. D02-E100/E300-05-N (batch volume ≥ 12 mL) or C02-E100-05-N (batch volume < 12 mL) filters with 300 kDa (HA and CMDex purifications) or 100 kDa (all other purifications) nominal molecular weight cutoffs were used to purify free dyes or polymers away from the nanoparticle samples. For PLR purification, the filter was pre-treated using a mock sample of free PLR, so saturate adsorption sites on the anionic membrane walls. Samples were filtered at 13 mL/min for small batches or 80 mL/min for large batches, with a Milli-Q water inlet line to replace 1:1 the volume of waste permeate. After at least 5 sample volume equivalents of waste collection, the sample was concentrated, removed from the filter by reversing the pump direction, and brought back to 1 mg/mL nanoparticle concentration (0.5 mg/mL for HA) by backflushing a defined volume of Milli-Q water through the filter and into the sample.

#### 2.5. Nanoparticle Characterization

Nanoparticle hydrodynamic size, polydispersity, and zeta potential were measured using dynamic light scattering (Malvern ZS90 Particle Analyzer, λ= 633 nm). 50 μg of each nanoparticle was diluted into 2 mM NaCl to give a total volume of 800 μL, then transferred to polystyrene cuvettes or DTS1070 folded capillary cuvettes for DLS.

### 3. Cell Culture

#### 3.1. Maintenance

hCMEC/D3 cells were cultured according to manufacturer specifications in EMB-2 media supplemented with 5% FBS, 1 ng/mL bFGF, 1.4 μM hydrocortisone, 1% pen/strep, 1% chemically defined lipid concentrate, 10 mM HEPES, and 5 μg/mL ascorbic acid. Cells were cultured in flasks coated with 12 μg/cm^2^ rat tail collagen and split twice per week at a ratio between 1:3 and 1:8, using Accumax dissociation reagent, to maintain cells below approximately 90% confluency. Between maintenance and experiments, cells were incubated at 37 °C and in a 100% humidity and 5% CO2 atmosphere. Cell lines were authenticated using STR profiling, and cells were tested monthly for mycoplasma, with all results coming back negative for contamination.

#### 3.2. Cell Monolayers for Gene Expression and Uptake Experiments

hCMEC/D3 cells were suspended in media and seeded at a density of 2 × 10^5^ cells/cm^2^ in collagen-coated (12 μg/cm^2^) transwell supports or 96-well tissue culture plates. The cells were incubated for 7 days, with media replacement every 2-3 days. For transwells, the TEER was monitored to confirm proper barrier formation before use in any experiments. TEER is expressed as the resistance of transwell filters with cells minus transwell filters without cells.

### 4. Transport Experiments

#### 4.1. General Protocol

Cell monolayers in transwells were transferred to 24-well plates containing 1 mL/well media. Nanoparticle or fluorescent marker treatments were added to the apical chambers at 20 μg/cm^2^ in media; negative control wells received fresh media. Extra treatments for each experiment were used to make fluorescence calibration curves. At 4, 8, and 24 hours after treatment addition, the monolayers were transferred to new basal plates (creating quasi-sink conditions for the basal media compartment). Media from the basal plates was sampled for fluorescence measurements on a Tecan Infinite M200 Pro plate reader, with 100 μL/well sample in a black 96-well plate, and applied to the calibration curves to calculate percent of nanomaterial transported.

#### 4.2. 4° C Protocol

To assess active transport, nanoparticle treatments in media and basal plates with media were both equilibrated to 4° C in the refrigerator. Pre-experiment TEER values were measured with cells still at 37° C, then monolayers were transferred to the chilled treatment and basal media. The monolayers were then incubated in the 4° C refrigerator for the remainder of the experiment.

#### 4.3. DMSO Breakup and Reread for Liposomes

To homogenize fluorescent liposome containing samples, 100 μL of sample (nanoparticles in cell culture media) per well was supplemented with 100 μL/well DMSO and 50 μL/well of heparin sulfate (1 mg/mL in PBS). The plate was placed on an orbital shaker at 240 RPM for 10 minutes before repeating the fluorescence measurements.

### 5. Gene Expression

#### 5.1. RNA extraction and cDNA preparation

Cells were rinsed with PBS and pelleted prior to RNA extraction. Total RNA was extracted according to the instructions provided with the Qiagen RNeasy kit. Briefly, lysis buffer was prepared with the recommended amount of β-mercaptoethanol to protect RNA from degradation. 1-3 million cells worth of lysate were added to spin columns. Total RNA was eluted from the columns using 30 μL nuclease-free water. Nanodrop (Thermo Fisher) spectrophotometry was used to assess RNA concentration and quality, and all 260/280 values were greater than 1.8. cDNA was synthesized according to manufacturer’s instructions using 1 μg of template RNA. cDNA was stored at −20° C or placed on ice for immediate use.

#### 5.2. Real time quantitative PCR

For qPCR reactions, cDNA was diluted 1:50 with nuclease-free water, and primers were diluted to 20x (10 μM) in IDTE buffer according to the manufacturer’s specifications. RT-qPCR was set up in a 384-well plate with 8 μL diluted cDNA, 10 μL 2x SYBR Green master mix, 0.8 μL nuclease-free water, and 1.2 μL 20x primer. Each condition was performed in technical triplicate. No primer (IDTE buffer instead of primer) and no cDNA (water instead of cDNA) controls were also used to ensure there was not contamination. RT-qPCR was run on a LightCycler 480 (Roche) and Ct values obtained using the second derivative. The ΔCt method was used to compare expression between cell lines, normalizing to beta actin. The primers have the following assay ID numbers and sequences:

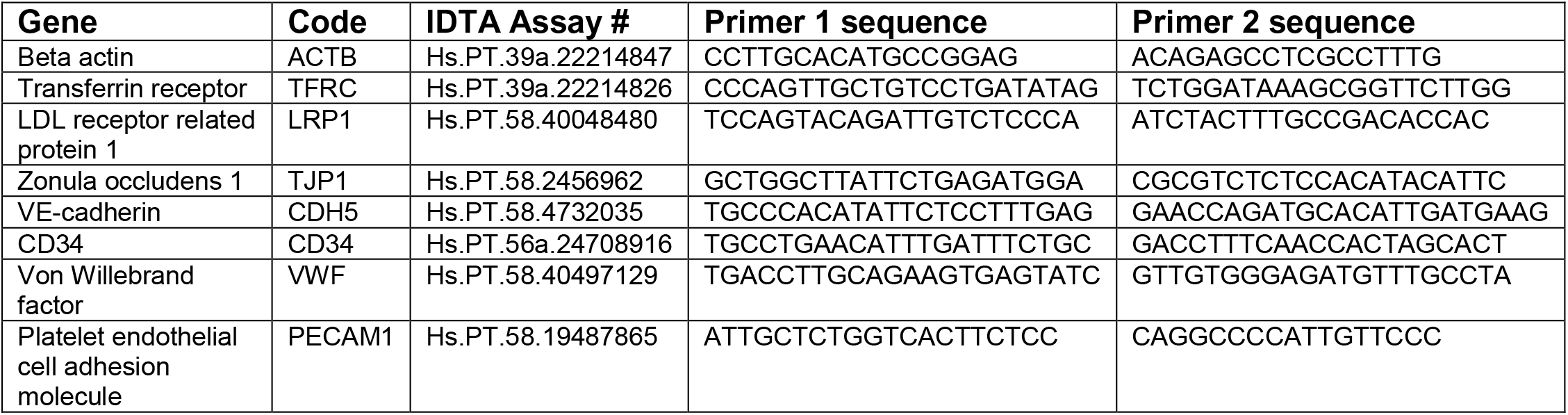

### 6. *in vitro* Confocal Imaging

#### 6.1. Sample treatment and preparation

Cells were cultured on collagen-coated chambered slides or transwells for seven days. The cells were incubated with Cy3-labelled NPs for 8 hours, then washed 3x with PBS and fixed with 4% formaldehyde in PBS for 20 minutes at room temperature, protected from light. This was followed by three washes with PBS at 5 min/wash, and samples were stored at 4° C overnight in fresh PBS. Samples were permeabilized and blocked with 0.1% saponin and 1% BSA in PBS for 30 minutes at room temperature. For samples with unlabeled primary antibodies, cells were stained for 30 minutes with primary solutions containing 0.1 % saponin, 1% BSA, and 2.5 μg/mL primary antibodies. The samples were rinsed three times with PBS over the course of fifteen minutes, then stained with solutions containing 2.5 μg/mL AlexaFluor 488 labelled antibodies, 1.5 μg/mL Hoechst 33342, and either 10 μg/mL wheat germ agglutinin or 0.165 μM phalloidin labelled with AlexaFluor 633. The samples were washed again with PBS three times over 15 minutes, then plastic chambers were removed from chamber slides, or transwell filters were punched out of their plastic casing using a biopsy punch and placed cell side up on a microscope slide. The samples were supplemented with Vectashield (H1000), secured with a coverslip, and sealed using nail polish. The samples were stored at 4 °C and protected from light until imaging.

#### 6.2. Image capture and processing

The cells were imaged with a confocal laser-scanning microscope (FV-1200, Olympus), equipped with 405, 473, 559, and 635 nm lasers. Images were acquired with 100x objectives, and all images were acquired under the same illumination settings, with the exception of samples treated with bare liposomes; these samples required higher laser settings and this caveat is noted in all of the relevant figure captions. Images were processed using ImageJ software. Bare liposome signal was linearly increased by 10%, and Syntaxin 6 background signal was reduced by raising the lower pixel limit by 10%. No other adjustments to image signals were made. With the exception of orthogonal views, images display Z projections (maximum signal) bounded by the center of the cell layer and the top of all four signals at the apical surface.

### 7. Flow Cytometry

Nanoparticle-treated cells in 96-well plates were washed 3x with warm PBS, then dissociated from the well bottoms using 30 μL Accumax dissociation reagent. 220 μL warm media was added to quench the dissociation, and pipetted vigorously to break up clumps before transferring to new 96-well plates without collagen coating. The samples were analyzed using a BD LSR II Flow Cytometer with a high throughput sampler (BD Biosciences). Samples dosed with Cy5 liposomes were analyzed on the APC channel (ex. 640, filters 670/30). Samples dosed with rhodamine B PLGA were analyzed on the PE channel (ex. 561 filters 582/42. Samples dosed with yellow green (polystyrene) fluospheres were analyzed on the GFP channel (ex. 488, filters 515/20). Data were analyzed using FlowJo (version 10), and cells were gated for single cells based on untreated cell samples using the side scatter and forward scatter plots.

### 8. Monolayer Association

Nanoparticle treated cells in 96-well plates were washed three times with ice cold PBS, then treated with 100 μL/well of 0.5 mg/mL heparin sulfate in 50% DMSO and 50% PBS. Standard curves were constructed by adding defined amounts of nanoparticles to the same homogenization solution. The samples were placed on an orbital shaker for 15 minutes at 180 RPM, then the samples transferred to black 96-well plates to measure particle fluorescence using the plate reader.

### 9. Animal Studies

#### 9.1. Animal Care and Use

All animal experiments were approved by the Massachusetts Institute of Technology Committee on Animal Care (CAC, protocol number 0919-056-22) and were conducted under the oversight of the Division of Comparative Medicine (DCM). C57BL/6 mice were purchased from Taconic, and were housed in cages of no more than five animals with controlled temperature (25 °C), 12 h light−dark cycles and free access to food and water. Both female and male mice were used in this study, and the mice were 8–20 weeks old at the time of experiment. The free-to-use PS power calculator (Vanderbilt) was used to determine the minimal sample size for which statistical power was greater than or equal to 0.8, leading to groups of 2 female + 2 male mice, with 1-3 measurements per animal.

#### 9.2. Intravital Imaging

Mice underwent head hair removal up to 24 hours before the imaging procedure occurred. One at a time, animals were injected with 70 kDa FITC-labelled dextran (2 mg/mL in PBS, sterile filtered) and red-fluorescent nanoparticles (1 mg/mL in 5% dextrose), both as 150 μL retro-orbital injections. To create the cranial window, the skull was exposed, and a high-speed hand drill (Dremel) was used to thin the skull until the dura mater was exposed over the right frontal cortex. The mice were then secured to a microscope coverslip for imaging. Multiphoton imaging was performed on an Olympus FV-1000MPE multiphoton microscope (Olympus) using a 25×, numerical aperture 1.05 objective. Excitation was achieved by using a femtosecond pulse laser at 840 nm, and emitted fluorescence was collected by photomultiplier tubes with emission filters of 425/30 nm for Collagen 1, 525/45 nm for fluorescein isothiocyanate-labeled dextran, 607/45 nm for red polystyrene nanoparticles, and 672/30 nm for Cy5 (PLGA and liposome) nanoparticles. Collagen 1 was excited by second harmonic generation and emitted as polarized light at half the excitation wavelength. The collagen 1 signal was used to identify the dura, such that the vessels imaged were within the cortex. ^19^ Images were acquired every 2 minutes for 12 minutes for analysis, and up to three image sessions per mouse were run. Mice were maintained under anesthesia for the duration of the imaging and then humanely euthanized.

Acquired images from intravital imaging were then thresholded and segmented by using the Fiji distribution of ImageJ and the Trainable Weka Segmentation plugin. Vessels below the dura and arteries were examined to ensure that these represent BBB capillaries in the mouse brain. The microvasculature filled with dextran (dextran channel) was used to generate a three-dimensional mask of the BBB mouse vessels. This mask was employed to calculate vessel surface area, as well as dextran and NP signal both inside and outside the blood vessels. After masking, analysis of NP or dextran transport was performed as previously described.^19,26,41^

### 10. Fluorescent polymer modifications

HA (60 kDa nominal molecular weight) and PSialA were dyed for intravital imaging using standard 1-Ethyl-3-[3-dimethylaminopropyl]carbodiimide (EDC) coupling with amine-functionalized SulfoCy5. Polymers, sulfo-N-hydroxysuccinimide (sulfo-NHS), sulfoCy5-amine, and EDC were added sequentially in a 1 monomer : 0.5 : 0.255 : 2.55 molar ratio for HA, or 1 monomer : 0.5 : 0.085 : 1.5 molar ratio for PSialA. Reactions were allowed to run overnight, then the polymers were purified away from free dye and reactants using Amicon Ultra-4 3 kDa spin filters, followed by dialysis in Milli-Q water using 3.5 kDa cutoff tubing. Dialysis continued until no dye could be detected in the dialysate.

### 11. Statistics and Data Availability

All data are available in the manuscript and accompanying supplementary tables.

Detailed statistical information is provided for each figure in the associated caption, and all tests used nonparametric comparisons based on sample sizes. Unless noted otherwise, for single comparisons, the Mann-Whitney test was used. For multiple comparison testing, the Kruskal-Wallis test was used.

## References

1. Feigin VL, Nichols E, Alam T, et al. Global, regional, and national burden of neurological disorders, 1990–2016: a systematic analysis for the Global Burden of Disease Study 2016. Lancet Neurol. 2019;18(5):459–480. doi:10.1016/S1474-4422(18)30499-X

2. Parodi A, Rudzińska M, Deviatkin AA, Soond SM, Baldin A V., Zamyatnin AA. Established and emerging strategies for drug delivery across the blood-brain barrier in brain cancer. Pharmaceutics. 2019;11(5). doi:10.3390/pharmaceutics11050245

3. Haumann R, Videira JC, Kaspers GJL, van Vuurden DG, Hulleman E. Overview of Current Drug Delivery Methods Across the Blood–Brain Barrier for the Treatment of Primary Brain Tumors. CNS Drugs. 2020;34(11):1121–1131. doi:10.1007/s40263-020-00766-w

4. Abbott NJ, Patabendige AAK, Dolman DEM, Yusof SR, Begley DJ. Structure and function of the blood-brain barrier. Neurobiol Dis. 2010;37(1):13–25. doi:10.1016/j.nbd.2009.07.030

5. Urich E, Lazic SE, Molnos J, Wells I, Freskgård PO. Transcriptional profiling of human brain endothelial cells reveals key properties crucial for predictive in vitro blood-brain barrier models. PLoS One. 2012;7(5). doi:10.1371/journal.pone.0038149

6. Hollmann EK, Bailey AK, Potharazu A V., Neely MD, Bowman AB, Lippmann ES. Accelerated differentiation of human induced pluripotent stem cells to blood-brain barrier endothelial cells. Fluids Barriers CNS. 2017;14(1):1–13. doi:10.1186/s12987-017-0059-0

7. Lu TM, Barcia Durán JG, Houghton S, Rafii S, Redmond D, Lis R. Human Induced Pluripotent Stem Cell-Derived Brain Endothelial Cells: Current Controversies. Front Physiol. 2021;12(March). doi:10.3389/fphys.2021.642812

8. Poller B, Gutmann H, Krähenbühl S, et al. The human brain endothelial cell line hCMEC/D3 as a human blood-brain barrier model for drug transport studies. J Neurochem. 2008;107(5):1358–1368. doi:10.1111/j.1471-4159.2008.05730.x

9. Eigenmann DE, Xue G, Kim KS, Moses A V., Hamburger M, Oufir M. Comparative study of four immortalized human brain capillary endothelial cell lines, hCMEC/D3, hBMEC, TY10, and BB19, and optimization of culture conditions, for an in vitro blood-brain barrier model for drug permeability studies. Fluids Barriers CNS. 2013;10(1). doi:10.1186/2045-8118-10-33

10. Weksler BB, Subileau EA, Perrière N, et al. Blood-brain barrier-specific properties of a human adult brain endothelial cell line. FASEB J. 2005;19(13):1872–1874. doi:10.1096/fj.04-3458fje

11. Daniels BP, Cruz-Orengo L, Pasieka TJ, et al. Immortalized human cerebral microvascular endothelial cells maintain the properties of primary cells in an in vitro model of immune migration across the blood brain barrier. J Neurosci Methods. 2013;212(1):173–179. doi:10.1016/j.jneumeth.2012.10.001

12. Ohtsuki S, Ikeda C, Uchida Y, et al. Quantitative targeted absolute proteomic analysis of transporters, receptors and junction proteins for validation of human cerebral microvascular endothelial cell line hCMEC/D3 as a human blood-brain barrier model. Mol Pharm. 2013;10(1):289–296. doi:10.1021/mp3004308

13. Dauchy S, Miller F, Couraud PO, et al. Expression and transcriptional regulation of ABC transporters and cytochromes P450 in hCMEC/D3 human cerebral microvascular endothelial cells. Biochem Pharmacol. 2009;77(5):897–909. doi:10.1016/j.bcp.2008.11.001

14. Rahman NA, Rasil AN ain HM, Meyding-Lamade U, et al. Immortalized endothelial cell lines for in vitro blood-brain barrier models: A systematic review. Brain Res. 2016;1642:532–545. doi:10.1016/j.brainres.2016.04.024

15. Toth AE, Nielsen SSE, Tomaka W, Abbott NJ, Nielsen MS. The endo-lysosomal system of bEnd.3 and hCMEC/D3 brain endothelial cells. Fluids Barriers CNS. 2019;16(1):1–13. doi:10.1186/s12987-019-0134-9

16. Hajal C, Le Roi B, Kamm RD, Maoz BM. Biology and Models of the Blood-Brain Barrier. Annu Rev Biomed Eng. 2021;23:359–384. doi:10.1146/annurev-bioeng-082120-042814

17. Brown TD, Nowak M, Bayles A V., et al. A microfluidic model of human brain (μHuB) for assessment of blood brain barrier. Bioeng Transl Med. 2019;4(2):1–13. doi:10.1002/btm2.10126

18. Campisi M, Shin Y, Osaki T, Hajal C, Chiono V, Kamm RD. 3D self-organized microvascular model of the human blood-brain barrier with endothelial cells, pericytes and astrocytes. Biomaterials. 2018;180:117–129. doi:10.1016/j.biomaterials.2018.07.014

19. Straehla JP, Hajal C, Safford HC, et al. A predictive microfluidic model of human glioblastoma to assess trafficking of blood-brain barrier-penetrant nanoparticles. Proc Natl Acad Sci. 2022;119(23):e2118697119. doi:10.1073/pnas.2118697119/-/DCSupplemental.Published

20. Schmid D, Buntz A, Hanh Phan TN, et al. Transcytosis of payloads that are non-covalently complexed to bispecific antibodies across the hCMEC/D3 blood-brain barrier model. Biol Chem. 2018;399(7):711–721. doi:10.1515/hsz-2017-0311

21. Gonzalez-Carter D, Goode AE, Fiammengo R, Dunlop IE, Dexter DT, Porter AE. Inhibition of Leptin– ObR Interaction Does not Prevent Leptin Translocation Across a Human Blood–Brain Barrier Model. J Neuroendocrinol. 2016;28(6). doi:10.1111/jne.12392

22. Papadia K, Markoutsa E, Antimisiaris SG. How do the physicochemical properties of nanoliposomes affect their interactions with the hCMEC/D3 cellular model of the BBB? Int J Pharm. 2016;509(1–2):431–438. doi:10.1016/j.ijpharm.2016.06.019

23. Wang C, Wu B, Wu Y, Song X, Zhang S, Liu Z. Camouflaging Nanoparticles with Brain Metastatic Tumor Cell Membranes: A New Strategy to Traverse Blood–Brain Barrier for Imaging and Therapy of Brain Tumors. Adv Funct Mater. 2020;30(14):1–11. doi:10.1002/adfm.201909369

24. Brown TD, Habibi N, Wu D, Lahann J, Mitragotri S. Effect of Nanoparticle Composition, Size, Shape, and Stiffness on Penetration across the Blood-Brain Barrier. ACS Biomater Sci Eng. 2020;6(9):4916–4928. doi:10.1021/acsbiomaterials.0c00743

25. Markoutsa E, Pampalakis G, Niarakis A, et al. Uptake and permeability studies of BBB-targeting immunoliposomes using the hCMEC/D3 cell line. Eur J Pharm Biopharm. 2011;77(2):265–274. doi:10.1016/j.ejpb.2010.11.015

26. Offeddu GS, Haase K, Gillrie MR, et al. An on-chip model of protein paracellular and transcellular permeability in the microcirculation. Biomaterials. 2019;212(May):115–125. doi:10.1016/j.biomaterials.2019.05.022

27. Srinivasan B, Kolli AR, Esch MB, Abaci HE, Shuler ML, Hickman JJ. TEER Measurement Techniques for In Vitro Barrier Model Systems. J Lab Autom. 2015;20(2):107–126. doi:10.1177/2211068214561025

28. Helms HC, Abbott NJ, Burek M, et al. In vitro models of the blood-brain barrier: An overview of commonly used brain endothelial cell culture models and guidelines for their use. J Cereb Blood Flow Metab. 2015;36(5):862–890. doi:10.1177/0271678X16630991

29. Muller AM, Hermanns MI, Skrzynski C, Nesslinger M, Muller K-M, Kirkpatrick CJ. Expression of the Endothelial Markers PECAM-1, vWf, and CD34 in Vivo and in Vitro. Exp Mol Pathol. 2002;72:221–229. doi:10.1006/exmp.2002.2424

30. Kalari KR, Thompson KJ, Nair AA, et al. BBBomics-human blood brain barrier transcriptomics hub. Front Neurosci. 2016;10(MAR):1–7. doi:10.3389/fnins.2016.00071

31. Zakharova M, Tibbe MP, Koch LS, et al. Transwell-Integrated 2 μ m Thick Transparent Polydimethylsiloxane Membranes with Controlled Pore Sizes and Distribution to Model the Blood-Brain Barrier. Adv Mater Technol. 2021;2100138(6). doi:10.1002/admt.202100138

32. Correa S, Choi KY, Dreaden EC, et al. Highly Scalable, Closed-Loop Synthesis of Drug-Loaded, Layer-by-Layer Nanoparticles. Adv Funct Mater. 2016;26(7):991–1003. doi:10.1002/adfm.201504385

33. Correa S, Boehnke N, Barberio AE, et al. Tuning Nanoparticle Interactions with Ovarian Cancer through Layer-by-Layer Modification of Surface Chemistry. ACS Nano. 2020;14(2):2224–2237. doi:10.1021/acsnano.9b09213

34. Wang M, Zhang Y, Feng J, et al. Preparation, characterization, and in vitro and in vivo investigation of chitosan-coated poly (d,l-lactide-co-glycolide) nanoparticles for intestinal delivery of exendin-4. Int J Nanomedicine. 2013;8:1141–1154. doi:10.2147/IJN.S41457

35. Ribovski L, de Jong E, Mergel O, et al. Low nanogel stiffness favors nanogel transcytosis across an in vitro blood–brain barrier. Nanomedicine Nanotechnology, Biol Med. 2021;34:102377. doi:10.1016/j.nano.2021.102377

36. Nowak M, Brown TD, Graham A, Helgeson ME, Mitragotri S. Size, shape, and flexibility influence nanoparticle transport across brain endothelium under flow. Bioeng Transl Med. 2019;(September):1–11. doi:10.1002/btm2.10153

37. Predescu SA, Predescu DN, Malik AB. Molecular determinants of endothelial transcytosis and their role in endothelial permeability. Am J Physiol - Lung Cell Mol Physiol. 2007;293(4). doi:10.1152/ajplung.00436.2006

38. Eskelinen EL. Roles of LAMP-1 and LAMP-2 in lysosome biogenesis and autophagy. Mol Aspects Med. 2006;27(5–6):495–502. doi:10.1016/j.mam.2006.08.005

39. Lam FC, Morton SW, Wyckoff J, et al. Enhanced efficacy of combined temozolomide and bromodomain inhibitor therapy for gliomas using targeted nanoparticles. Nat Commun. 2018;9(1). doi:10.1038/s41467-018-04315-4

40. DeOre BJ, Partyka PP, Fan F, Galie PA. CD44 mediates shear stress mechanotransduction in an in vitro blood-brain barrier model through small GTPases RhoA and Rac1. FASEB J. 2022;36(5):1–16. doi:10.1096/fj.202100822RR

41. Haase K, Gillrie MR, Hajal C, Kamm RD. Pericytes Contribute to Dysfunction in a Human 3D Model of Placental Microvasculature through VEGF-Ang-Tie2 Signaling. Adv Sci. 2019;6(23). doi:10.1002/advs.201900878

