## Supplementary Information for "Core material and surface chemistry of Layer-by-Layer (LbL) nanoparticles independently direct uptake, transport, and trafficking in preclinical blood-brain barrier (BBB) models"

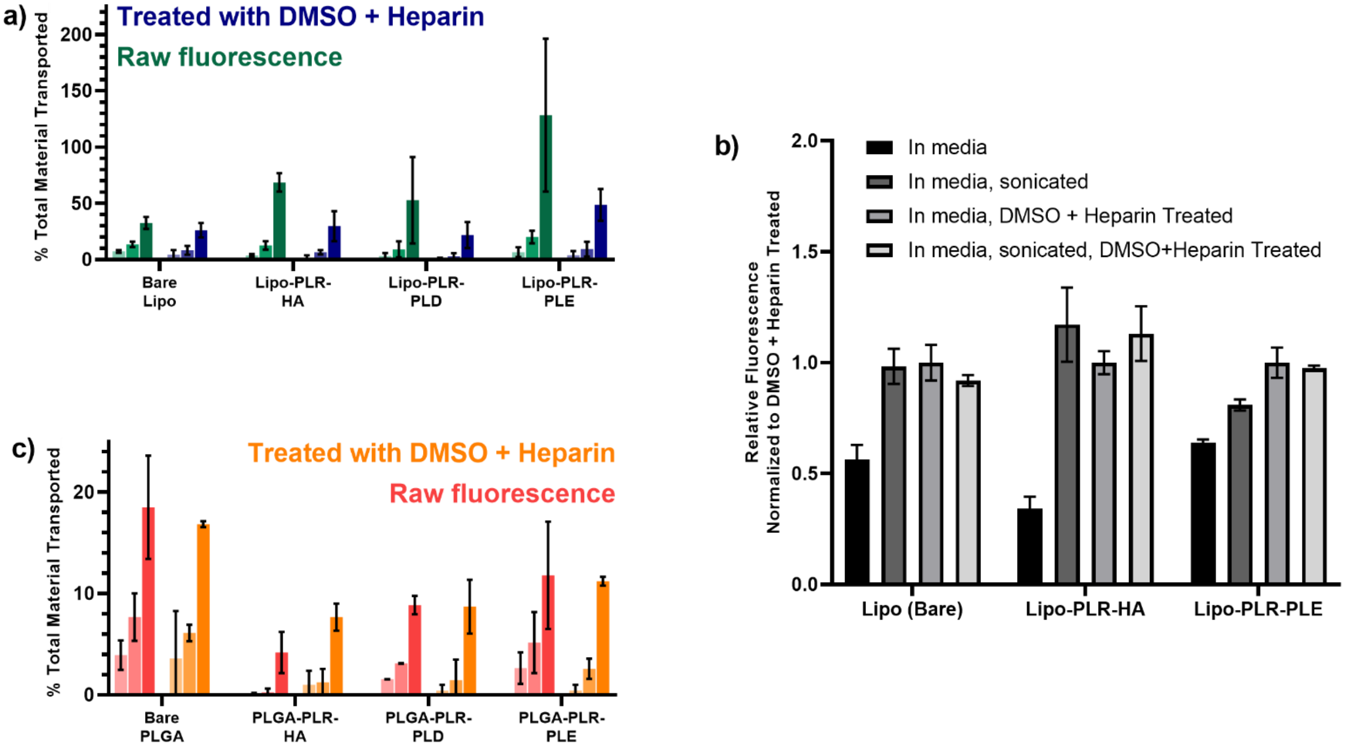


**Supplementary Figure 1: Liposome transport data in transwell models can be skewed by Cy5 fluorophore dequenching upon nanoparticle breakup. (a)** In a selection of liposome-based LbL-NPs, some outer layers – especially PLE – demonstrated drastic differences in transwell transport data depending on whether samples were treated with DMSO and heparin to break up NPs. **(b)** Nanoparticle fluorescent signal for all three formulations in cell culture media increased substantially and to approximately the same values after being broken up by sonicating, treating with DMSO + heparin sulfate, or both. Particle fluorescence data are normalized to samples treated with DMSO + heparin without sonication, and error displays standard deviation of 3 replicate wells of each sample. **(c)** By contrast, PLGA core LbL-NPs – which incorporate a rhodamine B fluorophore in quantities that do not self-quench – did not show these discrepancies. All transwell transport is reported as arithmetic mean ± standard deviation of three plate replicates, with three technical replicate wells per treatment per plate.

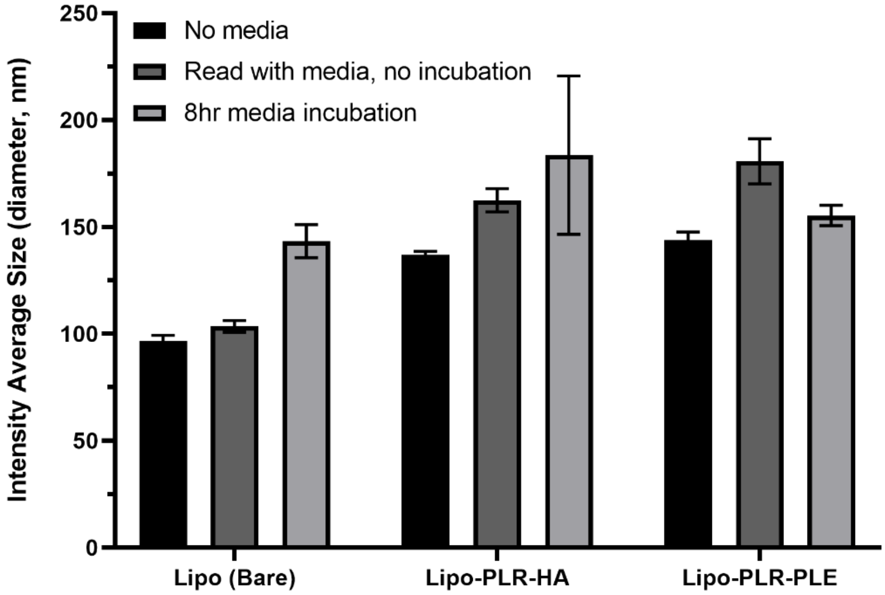


**Supplementary Figure 2: Nanoparticles do not substantially aggregate when incubated in cell culture media.** Nanoparticles were incubated in cell culture media for 8 hours at 37°C, then diluted into deionized water to measure size by DLS. Compared to control samples diluted in 2 mM NaCl or in 10% media (diluted in Milli-Q water) immediately before measuring, particles incubated in media displayed small changes in size consistent with protein adsorption but not particle aggregation. Data display mean ± standard deviation of three DLS measurements, and diameters are expressed as intensity average – rather than number average – as a metric that is more sensitive to smaller populations of particle aggregates.

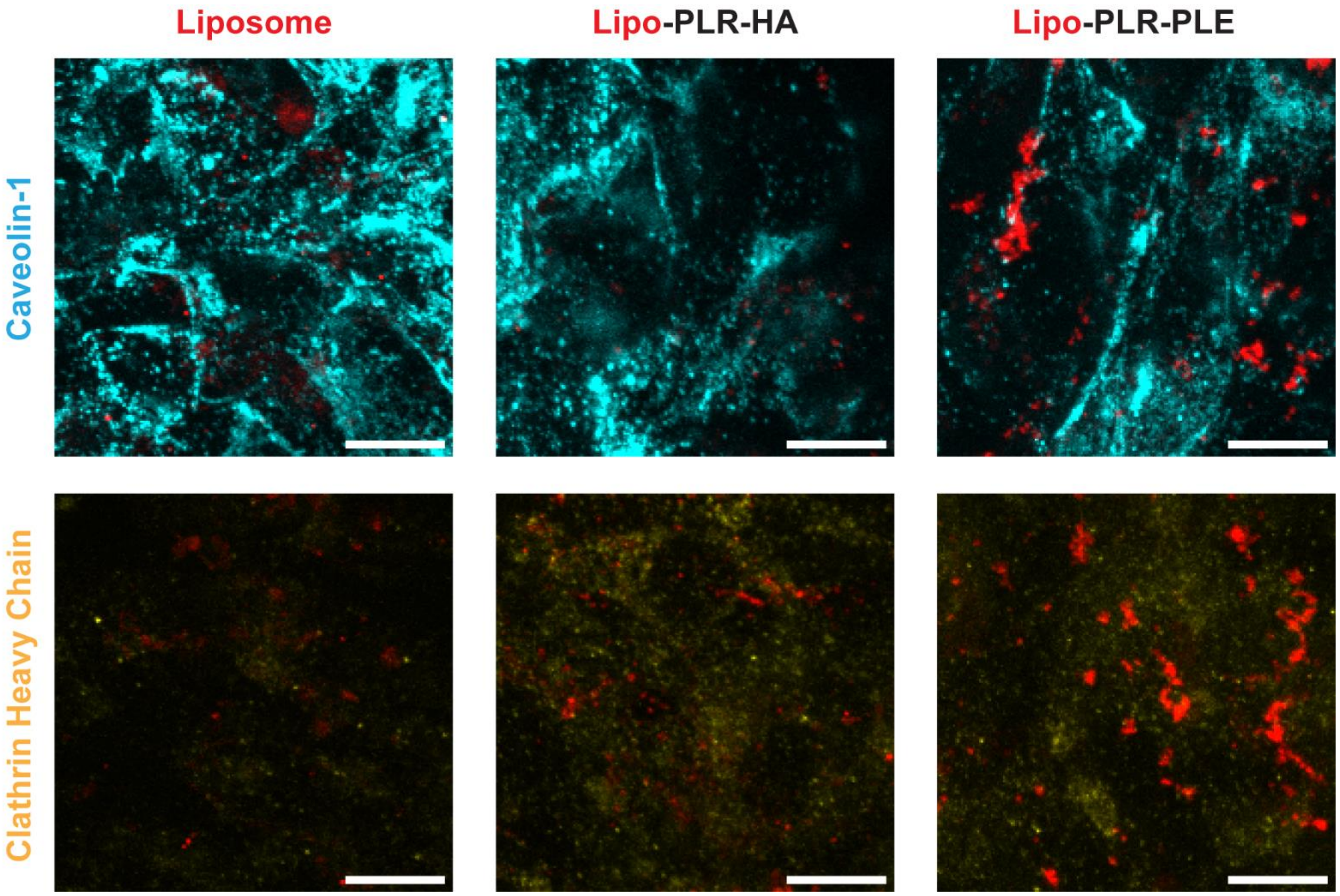


**Supplementary Figure 3: Nanoparticles did not appear to undergo uptake via caveolin- or clathrin-mediated endocytosis.** Nanoparticle signal for all three formulations did not colocalize with immunofluorescence signal for protein machinery for either uptake pathway. Scale bars display 10 µm. Bare liposomes required higher laser power for proper visualization, so quantity of nanoparticle signal should not be directly compared between particles.

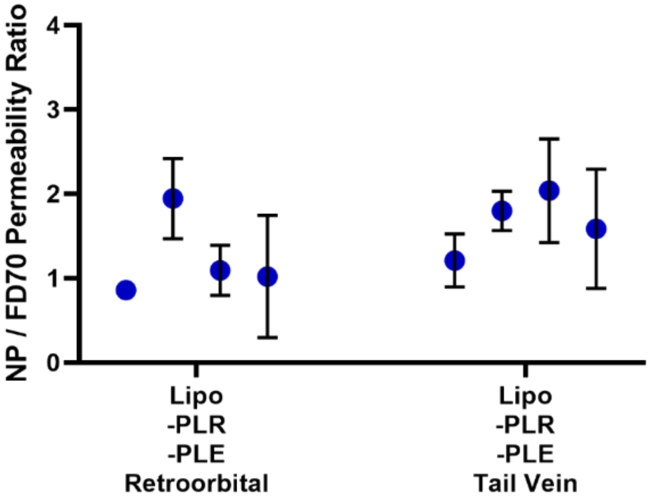


**Supplementary Figure 4: Retroorbital and tail vein injections yield comparable blood brain barrier permeability data for intravenously administered, fluorescent nanoparticles.** Data display mean ± s.e.m. for individual mice (n = 3-15 permeability measurements per mouse). Groups are not statistically significant at the $p < 0.05$ level by Mann-Whitney *U* test.

**Supplementary Table 1:** Gene expression values for transwell (Trans) and solid plasticware (Plastic) grown cells at given passage numbers (P#) and days of development (D#). All values are arithmetic mean of 3-4 well replicates.

| | P2D4 | | P2D7 | | P2D14 | | P6D4 | | P6D7 | | P6D14 | | P10D4 | | P10D7 | | P10D14 | |
|---|---|---|---|---|---|---|---|---|---|---|---|---|---|---|---|---|---|---|
| | Trans | Plastic | Trans | Plastic | Trans | Plastic | Trans | Plastic | Trans | Plastic | Trans | Plastic | Trans | Plastic | Trans | Plastic | Trans | Plastic |
| ACTB | 1.000 | 1.000 | 1.000 | 1.000 | 1.000 | 1.000 | 1.000 | 1.000 | 1.000 | 1.000 | 1.000 | 1.000 | 1.000 | 1.000 | 1.000 | 1.000 | 1.000 | 1.000 |
| TFRC | 0.021 | 0.020 | 0.092 | 0.100 | 0.034 | 0.041 | 0.063 | 0.083 | 0.022 | 0.024 | 0.017 | 0.017 | 0.024 | 0.037 | 0.057 | 0.045 | 0.015 | 0.016 |
| LRP1 | 0.048 | 0.037 | 0.034 | 0.044 | 0.077 | 0.065 | 0.025 | 0.027 | 0.027 | 0.018 | 0.042 | 0.035 | 0.020 | 0.012 | 0.030 | 0.017 | 0.048 | 0.041 |
| TJP1 | 0.233 | 0.155 | 0.103 | 0.180 | 0.235 | 0.291 | 0.049 | 0.058 | 0.113 | 0.074 | 0.111 | 0.118 | 0.154 | 0.088 | 0.239 | 0.242 | 0.165 | 0.133 |
| CDH5 | 0.026 | 0.013 | 0.050 | 0.045 | 0.035 | 0.037 | 0.030 | 0.028 | 0.023 | 0.026 | 0.018 | 0.011 | 0.008 | 0.006 | 0.027 | 0.030 | 0.009 | 0.007 |
| CD34 | 0.084 | 0.053 | 0.018 | 0.022 | 0.048 | 0.037 | 0.021 | 0.015 | 0.024 | 0.019 | 0.029 | 0.030 | 0.016 | 0.008 | 0.025 | 0.029 | 0.020 | 0.016 |
| VWF | 0.014 | 0.010 | 0.012 | 0.024 | 0.023 | 0.020 | 0.007 | 0.006 | 0.006 | 0.004 | 0.003 | 0.003 | 0.005 | 0.003 | 0.009 | 0.007 | 0.003 | 0.002 |
| PECAM1 | 0.049 | 0.034 | 0.012 | 0.012 | 0.107 | 0.055 | 0.009 | 0.008 | 0.020 | 0.011 | 0.018 | 0.014 | 0.014 | 0.007 | 0.019 | 0.025 | 0.014 | 0.022 |

| **Supplementary Table 2:** Nanoparticle characteristics for auxiliary nanoparticle batches, measured by dynamic light scattering. All values are displayed as mean ± SD of three runs. PDI: polydispersity index. | | | |
|---|---|---|---|
| Nanoparticle ID | Number Average Size (nm) | PDI | Zeta Potential (mV) |
| Cy3-labelled Nanoparticles for *in vitro* confocal imaging | | | |
| Bare Lipo | 74.5 ± 2.6 | 0.09 ± 0.00 | -46.7 ± 2.8 |
| Lipo-PLR-HA | 87.0 ± 6.3 | 0.14 ± 0.03 | -33.5 ± 0.7 |
| Lipo-PLR-PLE | 84.1 ± 2.8 | 0.15 ± 0.02 | -54.1 ± 1.9 |
| Cy5-labelled nanoparticles for intravital imaging in mice | | | |
| Bare Lipo | 70.8 ± 4.0 | 0.15 ± 0.04 | -42.0 ± 2.1 |
| Lipo-PLR-CMDex | 102.1 ± 13.8 | 0.23 ± 0.09 | -34.8 ± 1.0 |
| Lipo-PLR-HA | 105.7 ± 12.5 | 0.16 ± 0.04 | -31.2 ± 0.5 |
| Lipo-PLR-PLD | 102.2 ± 12.2 | 0.15 ± 0.00 | -39.9 ±1.9 |
| Lipo-PLR-PLE | 106.9 ± 3.9 | 0.17 ± 0.06 | -44.3 ± 3.1 |
| Lipo-PLR-PSialA | 107.0 ± 7.7 | 0.05 ± 0.04 | -38.3 ± 0.9 |
| PLGA-PLR-PLD | 134.4 ± 9.2 | 0.22 ± 0.05 | -43.0 ± 1.0 |
| PS-PLR-PLD | 132.4 ± 2.7 | 0.22 ± 0.09 | -46.4 ± 2.17 |

**Supplementary Table 3: Tabulated blood brain barrier permeability values for intravital imaging, displaying nanoparticle permeability (P NP), dextran permeability P FD70, and the ratio thereof for each animal.**

| | Imaging Session 1 | | | | | Imaging Session 2 | | | | | Imaging Session 3 | | | | |
|---|---|---|---|---|---|---|---|---|---|---|---|---|---|---|---|
| ID | P NP (µm/s) | P NP (cm/s) | P FD70 (µm/s) | P FD70 (cm/s) | P[NP]/ P[FD70] | P NP (µm/s) | P NP (cm/s) | P FD70 (µm/s) | P FD70 (cm/s) | P[NP]/ P[FD70] | P NP (µm/s) | P NP (cm/s) | P FD70 (µm/s) | P FD70 (cm/s) | P[NP]/ P[FD70] |
| | 4.00E-03 | 1.62E-03 | 1.62E-07 | 4.00E-07 | 2.47E+00 | | | | | | | | | | |
| | 1.62E-03 | 8.69E-04 | 8.69E-08 | 1.62E-07 | 1.86E+00 | | | | | | Did not take a third set of images | | | | |
| Lipo-1-F | 2.46E-04 | 2.88E-04 | 2.88E-08 | 2.46E-08 | 8.56E-01 | | | | | | | | | | |
| | Signals fade and permeability values become negative | | | | | | | | | | | | | | |
| | | | | | | | | | | | 2.05E-03 | 2.05E-07 | 4.39E-03 | 4.39E-07 | 4.68E-01 |
| | | | | | | | | | | | 2.82E-03 | 2.82E-07 | 2.44E-03 | 2.44E-07 | 1.16E+00 |
| Lipo-2-F | | | | | | | | | | | 3.30E-03 | 3.30E-07 | 1.47E-03 | 1.47E-07 | 2.25E+00 |
| | | | | | | | | | | | 2.99E-03 | 2.99E-07 | 1.09E-03 | 1.09E-07 | 2.73E+00 |
| | | | | | | | | | | | 1.64E-03 | 1.64E-07 | 9.36E-04 | 9.36E-08 | 1.75E+00 |
| | 2.55E-03 | 2.55E-07 | 1.92E-03 | 1.92E-07 | 1.33E+00 | | | | | | | | | | |
| | 1.32E-03 | 1.32E-07 | 1.44E-03 | 1.44E-07 | 9.20E-01 | | | | | | | | | | |
| Lipo-3-M | 7.30E-04 | 7.30E-08 | 1.14E-03 | 1.14E-07 | 6.42E-01 | | | | | | Did not take a third set of images | | | | |
| | 3.04E-04 | 3.04E-08 | 7.22E-04 | 7.22E-08 | 4.20E-01 | | | | | | | | | | |
| | 2.45E-04 | 2.45E-08 | 5.39E-04 | 5.39E-08 | 4.55E-01 | | | | | | | | | | |
| | | | | | | | | | | | 4.24E-03 | 4.24E-07 | 4.76E-03 | 4.76E-07 | 8.90E-01 |
| | | | | | | | | | | | 1.14E-02 | 1.14E-06 | 2.95E-03 | 2.95E-07 | 3.87E+00 |
| Lipo-4-M | Breathing artifacts interfere with image processing | | | | | Breathing artifacts interfere with image processing | | | | | 3.04E-03 | 3.04E-07 | 2.02E-03 | 2.02E-07 | 1.51E+00 |
| | | | | | | | | | | | 2.15E-03 | 2.15E-07 | 1.24E-03 | 1.24E-07 | 1.73E+00 |
| | | | | | | | | | | | 1.07E-03 | 1.07E-07 | 9.21E-04 | 9.21E-08 | 1.16E+00 |
| | 9.12E-03 | 9.12E-07 | 4.71E-03 | 4.71E-07 | 1.94E+00 | 5.24E-03 | 5.24E-07 | 3.20E-03 | 3.20E-07 | 1.64E+00 | Terminal complications early in this image set | | | | |
| HA-1-F | 1.71E-03 | 1.71E-07 | 1.61E-03 | 1.61E-07 | 1.06E+00 | 3.09E-03 | 3.09E-07 | 1.31E-03 | 1.31E-07 | 2.36E+00 | | | | | |
| | 1.78E-03 | 1.78E-07 | 1.36E-03 | 1.36E-07 | 1.31E+00 | Signals fade and permeability values become negative | | | | | | | | | |
| | 3.30E-03 | 3.30E-07 | 1.31E-03 | 1.31E-07 | 2.52E+00 | | | | | | | | | | |

| | | | | | | | | | | | | | | | |
|---|---|---|---|---|---|---|---|---|---|---|---|---|---|---|---|
| | 2.43E-03 | 2.43E-07 | 1.15E-03 | 1.15E-07 | 2.12E+00 | | | | | | | | | | |
| | 2.77E-02 | 2.77E-06 | 6.56E-03 | 3.00E+00 | 4.23E+00 | 4.60E-03 | 4.60E-07 | 2.76E-03 | 4.00E+00 | 1.67E+00 | 7.96E-03 | 7.96E-07 | 2.77E-03 | 2.77E-07 | 2.87E+00 |
| | 1.68E-02 | 1.68E-06 | 3.38E-03 | 3.38E-07 | 4.98E+00 | 3.27E-03 | 3.27E-07 | 1.44E-03 | 1.44E-07 | 2.28E+00 | 4.72E-03 | 4.72E-07 | 1.82E-03 | 1.82E-07 | 2.59E+00 |
| HA-2-F | 1.48E-02 | 1.48E-06 | 2.29E-03 | 2.29E-07 | 6.45E+00 | | | | | | 1.91E-03 | 1.91E-07 | 1.27E-03 | 1.27E-07 | 1.50E+00 |
| | 1.02E-02 | 1.02E-06 | | 2.11E-07 | 4.83E+00 | Signals fade and permeability values become negative | | | | | 1.75E-03 | 1.75E-07 | 8.96E-04 | 8.96E-08 | 1.95E+00 |
| | 9.63E-03 | 9.63E-07 | 1.90E-03 | 1.90E-07 | 5.08E+00 | | | | | | 1.36E-03 | 1.36E-07 | 7.53E-04 | 7.53E-08 | 1.80E+00 |
| | Signal jump - start calculations at second image | | | | | 3.52E-03 | 3.52E-07 | 6.86E-04 | 6.86E-08 | 5.13E+00 | 1.49E-03 | 1.49E-07 | 1.08E-03 | 1.08E-07 | 1.38E+00 |
| | 4.88E-03 | 4.88E-07 | 1.32E-03 | 1.32E-07 | 3.69E+00 | 5.77E-03 | 5.77E-07 | 3.37E-03 | 3.37E-07 | 1.71E+00 | 2.94E-03 | 2.94E-07 | 9.31E-04 | 9.31E-08 | 3.16E+00 |
| HA-3-M | 3.94E-03 | 3.94E-07 | 1.03E-03 | 1.03E-07 | 3.81E+00 | | | | | | | | | | |
| | 3.16E-03 | 3.16E-07 | 7.48E-04 | 7.48E-08 | 4.23E+00 | Signals fade and permeability values become negative | | | | | Signals fade and permeability values become negative | | | | |
| | 2.18E-03 | 2.18E-07 | 5.85E-04 | 5.85E-08 | 3.72E+00 | | | | | | | | | | |
| | 1.06E-03 | 1.06E-07 | 1.77E-03 | 1.77E-07 | 5.98E-01 | 6.22E-03 | 6.22E-07 | 1.96E-03 | 1.96E-07 | 2.30E+00 | 4.40E-03 | 4.40E-07 | 1.73E-03 | 1.73E-07 | 2.53E+00 |
| | 2.75E-03 | 2.75E-07 | 1.08E-03 | 1.08E-07 | 2.56E+00 | 3.94E-03 | 3.94E-07 | 1.06E-03 | 1.06E-07 | 3.71E+00 | 2.14E-03 | 2.14E-07 | 1.17E-03 | 1.17E-07 | 1.83E+00 |
| HA-4-M | 3.11E-03 | 3.11E-07 | 1.02E-03 | 1.02E-07 | 3.04E+00 | 3.34E-03 | 3.34E-07 | 9.02E-04 | 9.02E-08 | 5.12E+00 | 1.43E-03 | 1.43E-07 | 7.91E-04 | 7.91E-08 | 1.80E+00 |
| | Z drift / frame shift causes negative permeability values | | | | | 3.24E-03 | 3.24E-07 | 8.46E-04 | 8.46E-08 | 3.84E+00 | 4.48E-04 | 4.48E-08 | 7.60E-04 | 7.60E-08 | 5.90E-01 |
| | | | | | | 2.79E-03 | 2.79E-07 | 8.99E-04 | 8.99E-08 | 3.11E+00 | 2.76E-04 | 2.76E-08 | 3.65E-04 | 3.65E-08 | 7.57E-01 |
| | 1.13E-01 | 1.13E-05 | 1.14E-01 | 1.14E-05 | 9.86E-01 | | | | | | 1.57E-02 | 1.57E-06 | 2.34E-02 | 2.34E-06 | 6.70E-01 |
| | 8.35E-02 | 8.35E-06 | 1.08E-01 | 1.08E-05 | 7.70E-01 | | | | | | 6.71E-03 | 6.71E-07 | 1.09E-02 | 1.09E-06 | 6.16E-01 |
| PLE-1-F | 8.58E-02 | 8.58E-06 | 1.01E-01 | 1.01E-05 | 8.47E-01 | Signals fade immediately; permeability values all negative | | | | | 7.69E-03 | 7.69E-07 | 8.90E-03 | 8.90E-07 | 8.64E-01 |
| | 7.28E-02 | 7.28E-06 | 7.96E-02 | 7.96E-06 | 9.14E-01 | | | | | | 6.55E-03 | 6.55E-07 | 5.97E-03 | 5.97E-07 | 1.10E+00 |
| | 7.54E-02 | 7.54E-06 | 7.86E-02 | 7.86E-06 | 9.59E-01 | | | | | | 4.53E-03 | 4.53E-07 | 5.36E-03 | 5.36E-07 | 8.45E-01 |
| | | | | | | | | | | | 3.81E-03 | 3.81E-07 | 3.67E-03 | 3.67E-07 | 1.04E+00 |
| | | | | | | | | | | | 3.17E-03 | 3.17E-07 | 1.52E-03 | 1.52E-07 | 2.08E+00 |
| PLE-2-F | Massive drift renders images unprocessable | | | | | | | | | | 1.52E-03 | 1.52E-07 | 8.10E-04 | 8.10E-08 | 1.88E+00 |
| | | | | | | | | | | | 2.29E-03 | 2.29E-07 | 6.28E-04 | 6.28E-08 | 3.64E+00 |
| | | | | | | | | | | | 6.26E-04 | 6.26E-08 | 5.83E-04 | 5.83E-08 | 1.07E+00 |
| | | | | | | 4.10E-03 | 4.10E-07 | 2.64E-03 | 2.64E-07 | 1.56E+00 | | | | | |
| | | | | | | 7.10E-04 | 7.10E-08 | 1.32E-03 | 1.32E-07 | 5.39E-01 | | | | | |
| PLE-3-M | Massive drift renders images unprocessable | | | | | 1.26E-03 | 1.26E-07 | 1.07E-03 | 1.07E-07 | 1.18E+00 | | | | | |
| | | | | | | Signals fade and permeability values become negative | | | | | | | | | |

| | | | | | | | | | | | | | | | |
|---|---|---|---|---|---|---|---|---|---|---|---|---|---|---|---|
| PLE-4-M | | | | | | 5.75E-04 | 5.75E-08 | 2.33E-04 | 2.33E-08 | 2.47E+00 | Did not take a third set of images | | | | |
| | | | | | | 6.94E-04 | 6.94E-08 | 2.79E-04 | 2.79E-08 | 3.86E-01 | | | | | |
| | | | | | | 3.01E-04 | 3.01E-08 | 3.40E-04 | 3.40E-08 | 2.07E-01 | | | | | |
| | | | | | | 2.39E-04 | 2.39E-08 | 3.77E-04 | 3.77E-08 | 1.66E-01 | | | | | |
| | | | | | | Signals fade and permeability values become negative | | | | | | | | | |
| PLD-1-F | 1.25E-03 | 1.25E-07 | 3.75E-03 | 3.75E-07 | 3.32E-01 | | | | | | Terminal complications early in this image set | | | | |
| | 1.07E-03 | 1.07E-07 | 1.85E-03 | 1.85E-07 | 5.79E-01 | | | | | | | | | | |
| | 3.28E-03 | 3.28E-07 | 1.45E-03 | 1.45E-07 | 2.27E+00 | | | | | | | | | | |
| | 9.21E-04 | 9.21E-08 | 9.30E-04 | 9.30E-08 | 9.91E-01 | | | | | | | | | | |
| | 2.08E-03 | 2.08E-07 | 8.78E-04 | 8.78E-08 | 2.37E+00 | | | | | | | | | | |
| PLD-2-F | | | | | | 2.11E-03 | 2.11E-07 | 1.42E-03 | 1.42E-07 | 1.49E+00 | Terminal complications early in this image set | | | | |
| | | | | | | 2.32E-03 | 2.32E-07 | 6.84E-04 | 6.84E-08 | 3.39E+00 | | | | | |
| | | | | | | 3.82E-04 | 3.82E-08 | 7.48E-04 | 7.48E-08 | 5.11E-01 | | | | | |
| | | | | | | 8.85E-04 | 8.85E-08 | 6.70E-04 | 6.70E-08 | 1.32E+00 | | | | | |
| | | | | | | 2.05E-03 | 2.05E-07 | 7.07E-04 | 7.07E-08 | 2.89E+00 | | | | | |
| PLD-3-M | 1.02E-02 | 1.02E-06 | 5.48E-03 | 5.48E-07 | 1.87E+00 | Signals all fade over time and do not give processable data | | | | | Signals all fade over time and do not give processable data | | | | |
| | 3.67E-03 | 3.67E-07 | 2.43E-03 | 2.43E-07 | 1.51E+00 | | | | | | | | | | |
| | 4.00E-03 | 4.00E-07 | 1.61E-03 | 1.61E-07 | 2.48E+00 | | | | | | | | | | |
| | 2.59E-03 | 2.59E-07 | 1.29E-03 | 1.29E-07 | 2.01E+00 | | | | | | | | | | |
| | 7.77E-04 | 7.77E-08 | 9.88E-04 | 9.88E-08 | 7.86E-01 | | | | | | | | | | |
| PLD-4-M | 2.99E-03 | 2.99E-07 | 2.79E-03 | 2.79E-07 | 1.07E+00 | Too much background signal in both channels - interferes with permeability calculations | | | | | 1.30E-02 | 1.30E-06 | 4.38E-03 | 4.38E-07 | 2.96E+00 |
| | 1.38E-03 | 1.38E-07 | 1.45E-03 | 1.45E-07 | 9.54E-01 | | | | | | 5.45E-03 | 5.45E-07 | 2.18E-03 | 2.18E-07 | 2.49E+00 |
| | 4.44E-04 | 4.44E-08 | 5.84E-04 | 5.84E-08 | 7.61E-01 | | | | | | 3.62E-03 | 3.62E-07 | 1.80E-03 | 1.80E-07 | 2.01E+00 |
| | 3.23E-04 | 3.23E-08 | 4.80E-04 | 4.80E-08 | 6.73E-01 | | | | | | 9.20E-04 | 9.20E-08 | 1.47E-03 | 1.47E-07 | 6.28E-01 |
| | 3.55E-04 | 3.55E-08 | 5.23E-04 | 5.23E-08 | 6.79E-01 | | | | | | 1.21E-03 | 1.21E-07 | 1.11E-03 | 1.11E-07 | 1.09E+00 |
| PLGA-1-F | 1.73E-03 | 1.73E-07 | 8.33E-04 | 8.33E-08 | 2.08E+00 | 1.90E-03 | 1.90E-07 | 3.75E-03 | 3.75E-07 | 5.06E-01 | 2.15E-03 | 2.15E-07 | 3.91E-03 | 3.91E-07 | 5.49E-01 |
| | 8.05E-04 | 8.05E-08 | 3.76E-04 | 3.76E-08 | 2.14E+00 | 2.09E-03 | 2.09E-07 | 1.89E-03 | 1.89E-07 | 1.10E+00 | 2.75E-03 | 2.75E-07 | 2.15E-03 | 2.15E-07 | 1.28E+00 |
| | 5.04E-04 | 5.04E-08 | 5.69E-04 | 5.69E-08 | 8.86E-01 | 2.45E-03 | 2.45E-07 | 2.03E-03 | 2.03E-07 | 1.21E+00 | 2.47E-03 | 2.47E-07 | 1.47E-03 | 1.47E-07 | 1.68E+00 |
| | 4.91E-04 | 4.91E-08 | 7.49E-04 | 7.49E-08 | 6.56E-01 | Massive shift; ROI disappears | | | | | 1.90E-03 | 1.90E-07 | 1.29E-03 | 1.29E-07 | 1.47E+00 |
| | 3.62E-04 | 3.62E-08 | 9.88E-04 | 9.88E-08 | 3.67E-01 | | | | | | 1.63E-03 | 1.63E-07 | 1.23E-03 | 1.23E-07 | 1.32E+00 |

| | | | | | | | | | | | | | | | |
|---|---|---|---|---|---|---|---|---|---|---|---|---|---|---|---|
| | 1.44E-03 | 1.44E-07 | 1.68E-03 | 1.68E-07 | 8.54E-01 | 1.22E-03 | 1.22E-07 | 4.82E-03 | 4.82E-07 | 2.52E-01 | | | | | |
| | 1.35E-03 | 1.35E-07 | 1.67E-03 | 1.67E-07 | 8.06E-01 | 3.64E-03 | 3.64E-07 | 2.19E-03 | 2.19E-07 | 1.66E+00 | | | | | |
| PLGA-2-F | 1.04E-03 | 1.04E-07 | 1.54E-03 | 1.54E-07 | 6.72E-01 | 2.41E-03 | 2.41E-07 | 1.82E-03 | 1.82E-07 | 1.32E+00 | Did not take a third set of images | | | | |
| | 1.06E-03 | 1.06E-07 | 1.37E-03 | 1.37E-07 | 7.77E-01 | 1.91E-03 | 1.91E-07 | 1.43E-03 | 1.43E-07 | 1.34E+00 | | | | | |
| | 9.29E-04 | 9.29E-08 | 1.25E-03 | 1.25E-07 | 7.42E-01 | 1.41E-03 | 1.41E-07 | 1.18E-03 | 1.18E-07 | 1.20E+00 | | | | | |
| | | | | | | -9.81E-03 | -9.81E-07 | 6.43E-04 | 6.43E-08 | -1.53E+01 | 7.11E-04 | 7.11E-08 | 2.01E-03 | 2.01E-07 | 3.54E-01 |
| | | | | | | -2.38E-03 | -2.38E-07 | 6.03E-04 | 6.03E-08 | -3.94E+00 | -4.39E-05 | -4.39E-09 | 2.02E-03 | 2.02E-07 | -2.17E-02 |
| PLGA-3-M | Mid-frame Z shift renders image unprocessable | | | | | -2.04E-03 | -2.04E-07 | 3.64E-04 | 3.64E-08 | -5.60E+00 | -1.31E-03 | -1.31E-07 | 2.01E-03 | 2.01E-07 | -6.53E-01 |
| | | | | | | 8.01E-04 | 8.01E-08 | 7.11E-04 | 7.11E-08 | 1.13E+00 | -1.48E-03 | -1.48E-07 | 1.84E-03 | 1.84E-07 | -8.05E-01 |
| | | | | | | 2.10E-03 | 2.10E-07 | 7.01E-04 | 7.01E-08 | 3.00E+00 | -8.58E-04 | -8.58E-08 | 1.73E-03 | 1.73E-07 | -4.97E-01 |
| | 3.46E-03 | 3.46E-07 | 3.24E-03 | 3.24E-07 | 1.07E+00 | 2.13E-03 | 2.13E-07 | 2.50E-03 | 2.50E-07 | 8.53E-01 | -1.46E-03 | -1.46E-07 | 2.35E-03 | 2.35E-07 | -6.20E-01 |
| | 3.32E-03 | 3.32E-07 | 2.35E-03 | 2.35E-07 | 1.42E+00 | 1.97E-03 | 1.97E-07 | 2.28E-03 | 2.28E-07 | 8.65E-01 | -1.55E-04 | -1.55E-08 | 1.30E-03 | 1.30E-07 | -1.19E-01 |
| PLGA-4-M | 8.92E-04 | 8.92E-08 | 2.29E-03 | 2.29E-07 | 3.90E-01 | 2.61E-03 | 2.61E-07 | 1.40E-03 | 1.40E-07 | 1.86E+00 | 4.29E-05 | 4.29E-09 | 1.14E-03 | 1.14E-07 | 3.77E-02 |
| | 1.09E-03 | 1.09E-07 | 1.17E-03 | 1.17E-07 | 9.26E-01 | 2.32E-03 | 2.32E-07 | 1.27E-03 | 1.27E-07 | 1.82E+00 | 3.22E-04 | 3.22E-08 | 8.53E-04 | 8.53E-08 | 3.78E-01 |
| | 1.68E-03 | 1.68E-07 | 1.35E-03 | 1.35E-07 | 1.25E+00 | 2.48E-03 | 2.48E-07 | 1.01E-03 | 1.01E-07 | 2.46E+00 | 4.07E-04 | 4.07E-08 | 8.28E-04 | 8.28E-08 | 4.92E-01 |
| | 1.52E-03 | 1.52E-07 | 1.29E-03 | 1.29E-07 | 1.19E+00 | 1.77E-03 | 1.77E-07 | 1.93E-03 | 1.93E-07 | 9.14E-01 | 1.40E-03 | 1.40E-07 | 9.83E-04 | 9.83E-08 | 1.42E+00 |
| | 1.63E-03 | 1.63E-07 | 1.51E-03 | 1.51E-07 | 1.08E+00 | 1.00E-03 | 1.00E-07 | 1.08E-03 | 1.08E-07 | 9.23E-01 | 8.35E-04 | 8.35E-08 | 6.34E-04 | 6.34E-08 | 1.32E+00 |
| PS-1-F | 1.51E-03 | 1.51E-07 | 1.47E-03 | 1.47E-07 | 1.03E+00 | 1.18E-03 | 1.18E-07 | 1.31E-03 | 1.31E-07 | 9.05E-01 | 9.06E-04 | 9.06E-08 | 7.54E-04 | 7.54E-08 | 1.20E+00 |
| | 9.27E-04 | 9.27E-08 | 9.70E-04 | 9.70E-08 | 9.55E-01 | 1.35E-03 | 1.35E-07 | 1.38E-03 | 1.38E-07 | 9.78E-01 | 8.59E-04 | 8.59E-08 | 7.40E-04 | 7.40E-08 | 1.16E+00 |
| | 6.16E-04 | 6.16E-08 | 8.48E-04 | 8.48E-08 | 7.26E-01 | 1.24E-03 | 1.24E-07 | 1.03E-03 | 1.03E-07 | 1.20E+00 | 8.01E-04 | 8.01E-08 | 6.00E-04 | 6.00E-08 | 1.33E+00 |
| | 1.84E-03 | 1.84E-07 | 2.10E-03 | 2.10E-07 | 8.75E-01 | | | | | | | | | | |
| | 1.42E-03 | 1.42E-07 | 1.45E-03 | 1.45E-07 | 9.77E-01 | | | | | | | | | | |
| PS-2-F | 1.28E-03 | 1.28E-07 | 1.11E-03 | 1.11E-07 | 1.16E+00 | Terminal complications early in this image set | | | | | Did not take a third set of images | | | | |
| | 8.76E-04 | 8.76E-08 | 8.47E-04 | 8.47E-08 | 1.03E+00 | | | | | | | | | | |
| | 8.42E-04 | 8.42E-08 | 6.53E-04 | 6.53E-08 | 1.29E+00 | | | | | | | | | | |
| | 1.21E-03 | 1.21E-07 | 1.93E-03 | 1.93E-07 | 6.26E-01 | | | | | | | | | | |
| | 9.87E-04 | 9.87E-08 | 1.49E-03 | 1.49E-07 | 6.65E-01 | Did not take a second set of images | | | | | Did not take a third set of images | | | | |
| PS-3-M | 8.08E-04 | 8.08E-08 | 1.33E-03 | 1.33E-07 | 6.08E-01 | | | | | | | | | | |
| | Terminal complications | | | | | | | | | | | | | | |

| | | | | | | | | | | | | | | | |
|---|---|---|---|---|---|---|---|---|---|---|---|---|---|---|---|
| | 3.05E-03 | 3.05E-07 | 2.97E-03 | 2.97E-07 | 1.02E+00 | 3.76E-03 | 3.76E-07 | 3.86E-03 | 3.86E-07 | 9.74E-01 | 2.44E-03 | 2.44E-07 | 2.85E-03 | 2.85E-07 | 8.54E-01 |
| | 1.99E-03 | 1.99E-07 | 2.07E-03 | 2.07E-07 | 9.59E-01 | 1.49E-03 | 1.49E-07 | 1.48E-03 | 1.48E-07 | 1.00E+00 | 1.51E-03 | 1.51E-07 | 1.80E-03 | 1.80E-07 | 8.40E-01 |
| PS-4-M | 1.86E-03 | 1.86E-07 | 1.87E-03 | 1.87E-07 | 9.91E-01 | 1.67E-03 | 1.67E-07 | 1.65E-03 | 1.65E-07 | 1.01E+00 | 1.16E-03 | 1.16E-07 | 1.45E-03 | 1.45E-07 | 7.98E-01 |
| | 4.65E-04 | 4.65E-08 | 5.13E-04 | 5.13E-08 | 9.06E-01 | 7.31E-04 | 7.31E-08 | 7.36E-04 | 7.36E-08 | 9.93E-01 | 1.12E-03 | 1.12E-07 | 1.44E-03 | 1.44E-07 | 7.77E-01 |
| | 4.65E-04 | 4.65E-08 | 4.88E-04 | 4.88E-08 | 9.53E-01 | 5.75E-04 | 5.75E-08 | 6.20E-04 | 6.20E-08 | 9.28E-01 | 9.83E-04 | 9.83E-08 | 1.25E-03 | 1.25E-07 | 7.88E-01 |
| | 8.50E-04 | 8.50E-08 | 1.54E-03 | 1.54E-07 | 5.53E-01 | | | | | | | | | | |
| | 1.97E-03 | 1.97E-07 | 1.80E-03 | 1.80E-07 | 1.10E+00 | | | | | | | | | | |
| PLE.TV-1-F | 2.38E-03 | 2.38E-07 | 1.36E-03 | 1.36E-07 | 1.75E+00 | | | | | | | | | | |
| | 3.53E-04 | 3.53E-08 | 6.47E-04 | 6.47E-08 | 5.46E-01 | | | | | | | | | | |
| | 1.58E-03 | 1.58E-07 | 7.51E-04 | 7.51E-08 | 2.11E+00 | | | | | | | | | | |
| | | | | | | 1.10E-03 | 1.10E-07 | 1.12E-03 | 1.12E-07 | 9.79E-01 | | | | | |
| | | | | | | 1.11E-03 | 1.11E-07 | 6.50E-04 | 6.50E-08 | 1.71E+00 | | | | | |
| PLE.TV-2-F | | | | | | 7.22E-04 | 7.22E-08 | 3.35E-04 | 3.35E-08 | 2.16E+00 | | | | | |
| | | | | | | 6.65E-04 | 6.65E-08 | 2.87E-04 | 2.87E-08 | 2.32E+00 | | | | | |
| | | | | | | 5.29E-04 | 5.29E-08 | 2.89E-04 | 2.89E-08 | 1.83E+00 | | | | | |
| | | | | | | | | | | | 1.16E-02 | 1.16E-06 | 3.21E-03 | 3.21E-07 | 3.63E+00 |
| | | | | | | | | | | | 5.31E-03 | 5.31E-07 | 1.64E-03 | 1.64E-07 | 3.24E+00 |
| PLE.TV-3-M | | | | | | | | | | | 1.30E-03 | 1.30E-07 | 6.85E-04 | 6.85E-08 | 1.90E+00 |
| | | | | | | | | | | | 3.49E-04 | 3.49E-08 | 5.17E-04 | 5.17E-08 | 6.74E-01 |
| | | | | | | | | | | | 3.21E-04 | 3.21E-08 | 4.34E-04 | 4.34E-08 | 7.39E-01 |
| | | | | | | | | | | | 2.62E-03 | 2.62E-07 | 3.24E-03 | 3.24E-07 | 8.09E-01 |
| | | | | | | | | | | | 4.42E-03 | 4.42E-07 | 1.32E-03 | 1.32E-07 | 3.35E+00 |
| PLE.TV-4-M | | | | | | Lost water on objective ; signals disappear | | | | | 2.50E-03 | 2.50E-07 | 1.23E-03 | 1.23E-07 | 2.02E+00 |
| | | | | | | | | | | | 1.40E-04 | 1.40E-08 | 9.40E-04 | 9.40E-08 | 1.49E-01 |
| | | | | | | | | | | | Signals fade and permeability values become negative | | | | |
| | | | | | | 3.03E-03 | 3.03E-07 | 1.02E-03 | 1.02E-07 | 2.96E+00 | | | | | |
| | | | | | | 1.48E-03 | 1.48E-07 | 5.93E-04 | 5.93E-08 | 2.50E+00 | | | | | |
| CMDex-1-F | | | | | | 4.31E-04 | 4.31E-08 | 3.24E-04 | 3.24E-08 | 1.33E+00 | | | | | |
| | | | | | | 5.40E-04 | 5.40E-08 | 2.75E-04 | 2.75E-08 | 1.96E+00 | | | | | |
| | | | | | | 4.31E-04 | 4.31E-08 | 3.19E-04 | 3.19E-08 | 1.35E+00 | | | | | |

| | | | | | | | | | | | | | | | |
|---|---|---|---|---|---|---|---|---|---|---|---|---|---|---|---|
| CMDex-2-F | | | | | | | | | | | 1.80E-03 | 1.80E-07 | 1.34E-03 | 1.34E-07 | 1.34E+00 |
| | | | | | | | | | | | 2.17E-03 | 2.17E-07 | 9.00E-04 | 9.00E-08 | 2.41E+00 |
| | | | | | | | | | | | 2.33E-03 | 2.33E-07 | 7.52E-04 | 7.52E-08 | 3.10E+00 |
| | | | | | | | | | | | 1.99E-03 | 1.99E-07 | 8.64E-04 | 8.64E-08 | 2.30E+00 |
| | | | | | | | | | | | Z drift / frame shift causes misalignment with mask | | | | |
| CMDex-3-M | | | | | | | | | | | 1.97E-03 | 1.97E-07 | 9.24E-04 | 9.24E-08 | 2.13E+00 |
| | | | | | | | | | | | 3.41E-04 | 3.41E-08 | 7.13E-04 | 7.13E-08 | 4.79E-01 |
| | | | | | | | | | | | 2.05E-04 | 2.05E-08 | 7.49E-04 | 7.49E-08 | 2.74E-01 |
| | | | | | | | | | | | 5.77E-04 | 5.77E-08 | 8.35E-04 | 8.35E-08 | 6.91E-01 |
| | | | | | | | | | | | 1.20E-03 | 1.20E-07 | 7.23E-04 | 7.23E-08 | 1.66E+00 |
| CMDex-4-M | 1.18E-02 | 1.18E-06 | 5.32E-03 | 5.32E-07 | 2.22E+00 | 5.34E-04 | 5.34E-08 | 9.50E-04 | 9.50E-08 | 5.62E-01 | Did not take a third set of images | | | | |
| | 8.03E-03 | 8.03E-07 | 2.83E-03 | 2.83E-07 | 2.84E+00 | 2.74E-03 | 2.74E-07 | 7.33E-04 | 7.33E-08 | 3.73E+00 | | | | | |
| | Signals fade and permeability values become negative | | | | | Signals fade and permeability values become negative | | | | | | | | | |
| FreeHA-1-F | 3.94E-03 | 3.94E-07 | 3.80E-03 | 3.80E-07 | 1.04E+00 | | | | | | | | | | |
| | 1.81E-03 | 1.81E-07 | 1.62E-03 | 1.62E-07 | 1.11E+00 | | | | | | | | | | |
| | 1.14E-03 | 1.14E-07 | 1.03E-03 | 1.03E-07 | 1.11E+00 | | | | | | | | | | |
| | 8.09E-04 | 8.09E-08 | 6.94E-04 | 6.94E-08 | 1.17E+00 | | | | | | | | | | |
| | 6.44E-04 | 6.44E-08 | 5.39E-04 | 5.39E-08 | 1.20E+00 | | | | | | | | | | |
| FreeHA-2-F | | | | | | 1.58E-03 | 1.58E-07 | 2.38E-03 | 2.38E-07 | 6.64E-01 | Did not take a third set of images | | | | |
| | | | | | | 1.58E-03 | 1.58E-07 | 2.22E-03 | 2.22E-07 | 7.14E-01 | | | | | |
| | | | | | | 1.72E-03 | 1.72E-07 | 2.36E-03 | 2.36E-07 | 7.26E-01 | | | | | |
| | | | | | | 1.11E-03 | 1.11E-07 | 1.20E-03 | 1.20E-07 | 9.22E-01 | | | | | |
| | | | | | | 1.21E-03 | 1.21E-07 | 1.21E-03 | 1.21E-07 | 9.94E-01 | | | | | |
| FreeHA-3-M | 2.12E-03 | 2.12E-07 | 1.26E-03 | 1.26E-07 | 1.68E+00 | | | | | | Did not take a third set of images | | | | |
| | 1.66E-03 | 1.66E-07 | 1.22E-03 | 1.22E-07 | 1.36E+00 | | | | | | | | | | |
| | 1.13E-03 | 1.13E-07 | 1.02E-03 | 1.02E-07 | 1.11E+00 | | | | | | | | | | |
| | 5.79E-04 | 5.79E-08 | 7.37E-04 | 7.37E-08 | 7.86E-01 | | | | | | | | | | |
| | 8.70E-04 | 8.70E-08 | 9.42E-04 | 9.42E-08 | 9.24E-01 | | | | | | | | | | |

| | | | | | | | | | | | | | | | |
|---|---|---|---|---|---|---|---|---|---|---|---|---|---|---|---|
| | | | | | | | | | | | 3.72E-03 | 3.72E-07 | 4.47E-03 | 4.47E-07 | 8.32E-01 |
| | | | | | | | | | | | 1.88E-03 | 1.88E-07 | 2.49E-03 | 2.49E-07 | 7.57E-01 |
| FreeHA-4-M | | | | | | | | | | | 1.03E-03 | 1.03E-07 | 1.35E-03 | 1.35E-07 | 7.65E-01 |
| | | | | | | | | | | | 6.63E-04 | 6.63E-08 | 9.94E-04 | 9.94E-08 | 6.67E-01 |
| | | | | | | | | | | | 6.71E-04 | 6.71E-08 | 8.80E-04 | 8.80E-08 | 7.63E-01 |
| | | | | | | 5.46E-03 | 5.46E-07 | 2.37E-03 | 2.37E-07 | 2.30E+00 | | | | | |
| | | | | | | 3.56E-03 | 3.56E-07 | 1.40E-03 | 1.40E-07 | 2.54E+00 | | | | | |
| PSialA-1-F | | | | | | 2.21E-03 | 2.21E-07 | 8.37E-04 | 8.37E-08 | 2.65E+00 | | | | | |
| | | | | | | 3.02E-03 | 3.02E-07 | 6.87E-04 | 6.87E-08 | 4.40E+00 | | | | | |
| | | | | | | 1.80E-03 | 1.80E-07 | 6.85E-04 | 6.85E-08 | 2.63E+00 | | | | | |
| | | | | | | 4.97E-03 | 4.97E-07 | 2.55E-03 | 2.55E-07 | 1.95E+00 | | | | | |
| | | | | | | 3.13E-03 | 3.13E-07 | 1.42E-03 | 1.42E-07 | 2.21E+00 | | | | | |
| PSialA-2-F | | | | | | 2.80E-03 | 2.80E-07 | 9.66E-04 | 9.66E-08 | 2.90E+00 | | | | | |
| | | | | | | 2.08E-03 | 2.08E-07 | 7.14E-04 | 7.14E-08 | 2.91E+00 | | | | | |
| | | | | | | 2.16E-03 | 2.16E-07 | 8.21E-04 | 8.21E-08 | 2.63E+00 | | | | | |
| | | | | | | 1.00E-03 | 1.00E-07 | 1.23E-03 | 1.23E-07 | 8.13E-01 | | | | | |
| | | | | | | 7.80E-04 | 7.80E-08 | 7.30E-04 | 7.30E-08 | 1.07E+00 | | | | | |
| PSialA-3-M | | | | | | 7.00E-04 | 7.00E-08 | 5.85E-04 | 5.85E-08 | 1.20E+00 | | | | | |
| | | | | | | 5.59E-04 | 5.59E-08 | 5.42E-04 | 5.42E-08 | 1.03E+00 | | | | | |
| | | | | | | 5.27E-04 | 5.27E-08 | 4.85E-04 | 4.85E-08 | 1.09E+00 | | | | | |
| | 1.05E-02 | 1.05E-06 | 5.36E-03 | 5.36E-07 | 1.95E+00 | | | | | | | | | | |
| | 6.06E-03 | 6.06E-07 | 3.91E-03 | 3.91E-07 | 1.55E+00 | | | | | | | | | | |
| PSialA-4-M | 5.25E-03 | 5.25E-07 | 3.79E-03 | 3.79E-07 | 1.38E+00 | | | | | | Did not take a third set of images | | | | |
| | 4.19E-03 | 4.19E-07 | 2.54E-03 | 2.54E-07 | 1.65E+00 | | | | | | | | | | |
| | 3.45E-03 | 3.45E-07 | 2.22E-03 | 2.22E-07 | 1.55E+00 | | | | | | | | | | |
| | | | | | | | | | | | 1.67E-03 | 1.67E-07 | 1.94E-03 | 1.94E-07 | 8.57E-01 |
| | | | | | | | | | | | 1.79E-03 | 1.79E-07 | 1.56E-03 | 1.56E-07 | 1.14E+00 |
| FreePSialA-1-F | | | | | | | | | | | 1.08E-03 | 1.08E-07 | 1.26E-03 | 1.26E-07 | 8.63E-01 |
| | | | | | | | | | | | 7.41E-04 | 7.41E-08 | 9.88E-04 | 9.88E-08 | 7.50E-01 |
| | | | | | | | | | | | 3.78E-04 | 3.78E-08 | 9.05E-04 | 9.05E-08 | 4.18E-01 |

| | | | | | | | | | | |
|---|---|---|---|---|---|---|---|---|---|---|
| | | | | | | 3.46E-03 | 3.46E-07 | 1.16E-03 | 1.16E-07 | 2.97E+00 |
| | | | | | | Signals temporarily dip and permeability goes negative | | | | |
| FreePSialA-2-F | | | | | | 1.33E-04 | 1.33E-08 | 3.36E-04 | 3.36E-08 | 3.95E-01 |
| | | | | | | 3.58E-04 | 3.58E-08 | 2.89E-04 | 2.89E-08 | 1.24E+00 |
| | | | | | | 2.79E-04 | 2.79E-08 | 1.88E-04 | 1.88E-08 | 1.48E+00 |
| | | | | | | 5.75E-03 | 5.75E-07 | 2.84E-03 | 2.84E-07 | 2.03E+00 |
| | | | | | | 1.23E-03 | 1.23E-07 | 2.01E-03 | 2.01E-07 | 6.09E-01 |
| FreePSialA-3-M | | | | | | Z shifts out of frame and gives nonsense data for image 3-->4 and 4--> 5, but is salvageable for 5-->6 | | | | |
| | | | | | | 2.01E-03 | 2.01E-07 | 1.44E-03 | 1.44E-07 | 1.40E+00 |
| | 1.98E-03 | 1.98E-07 | 2.73E-03 | 2.73E-07 | 7.26E-01 | | | | | |
| | 1.91E-03 | 1.91E-07 | 1.69E-03 | 1.69E-07 | 1.13E+00 | | | | | |
| FreePSialA-4-M | 3.52E-03 | 3.52E-07 | 2.01E-03 | 2.01E-07 | 1.75E+00 | Not enough vessels to be quantifiable | | | | |
| | 2.52E-03 | 2.52E-07 | 1.66E-03 | 1.66E-07 | 1.52E+00 | | | | | |
| | 2.02E-03 | 2.02E-07 | 1.35E-03 | 1.35E-07 | 1.50E+00 | | | | | |
